# GlialCAM Cytoplasmic Signaling in Oligodendrocytes and Astrocytes is Essential for White Matter Homeostasis in the Brain

**DOI:** 10.1101/2025.10.03.680331

**Authors:** Zahra Nassiri Toosi, Santiago A. Forero, Ali Pirani, Sumod Sebastian, Tao Wang, Zhihua Chen, Xiaofeng Zheng, John E. Morales, Joseph H. McCarty

## Abstract

Glial cell adhesion molecule (GlialCAM) is an astrocyte- and oligodendrocyte-expressed transmembrane protein with two extracellular IgG-like domains and a cytoplasmic tail with putative signaling functions. While numerous studies have explored functions for the GlialCAM IgG-like domains in brain development and physiology, functions for its cytoplasmic signaling tail remain largely unknown. Therefore, we developed a mutant mouse model that expresses a truncated GlialCAM construct (GlialCAM ΔCT) that contains intact extracellular and transmembrane domains but lacks the cytoplasmic tail. Deletion of the GlialCAM cytoplasmic domain in glial cells of the brain results in vacuolization within white matter regions without disrupting neurovascular barrier integrity. Consequently, mutant mice exhibited selective deficits in motor coordination, muscular strength, and memory. Single cell transcriptome sequencing identifies GlialCAM-dependent defects in ECM remodeling pathways in white matter tracts. In situ spatial profiling revealed robust activation of astrocytes and microglia in the mutant brain. Proteomic analysis identified GlialCAM cytoplasmic tail interactors with links to MAPK signaling and cytoskeletal regulatory networks. These data reveal important functions for the GlialCAM cytoplasmic tail in homeostasis of white matter tracts in the adult murine brain. The GlialCAM ΔCT model may also be useful for studying the pathogenesis and possible treatment of neurological diseases linked to white matter degeneration.

## INTRODUCTION

GlialCAM, encoded by the *Hepacam* gene, is a 418 amino acid protein consisting of two extracellular Ig-like domains, a single pass transmembrane domain, and a cytoplasmic tail containing several putative signaling motifs ^1^. Hepacam was originally cloned from human liver cells and was subsequently found to be enriched in glial cells of the central nervous system (CNS)^2^. GlialCAM is predominantly expressed in white matter regions of the brain, with its expression profile closely paralleling that of myelin basic protein (MBP), as both show a temporal increase during postnatal brain development in mice ^2^. While GlialCAM is particularly enriched in perivascular astrocytes ^3^, it’s also expressed in oligodendrocytes and at low levels in some neurons ^2–5^. Additionally, GlialCAM is present in astroglial-derived exosomes, suggesting roles in axon outgrowth and/or dendritic spine formation ^6^. Heritable mutations in HEPACAM are linked to the pathogenesis of MLC, a rare disorder characterized by subcortical cysts and white matter edema ^3,7^. Most mutations in HEPACAM are mapped to the extracellular IgG-like domains, which are critical for protein localization to cell-cell junctions. These mutations disrupt the proper targeting of ion-water homeostasis mediators, ClC-2 and MLC1, to cell-cell junctions ^3,8^. GlialCAM not only affects the sub cellular localization of ClC-2, but also modulates its activity through its N-terminal extracellular regions ^8^. Mutant mouse models in which these genes have been targeted develop progressive myelin vacuolization, highlighting the importance of these membrane proteins in white matter integrity ^9^.

GlialCAM also interacts with the gap junction protein connexin 43 (Cx43) to maintain ion and water hemostasis in white matter regions of the brain ^10^. GlialCAM is also required for establishing boundaries between neighboring astrocytes and organizing gap junction coupling between astrocytes. These functions are critical for shaping astrocyte morphology during brain development ^4^. Recent proteomics studies have identified GlialCAM-interacting proteins, revealing functional associations with G protein-coupled receptors such as GPRC5B and GPR37L1, which are involved in modulating astrocyte volume regulation and maturation. Disrupting GlialCAM interactions with these partners results in osmotic pressure and ion gradient imbalances potentially linked to blood vessels, leading to astrocyte swelling and formation of vacuoles within myelin sheaths ^11–13^. Whether the GlialCAM extracellular IgG-like domains or intracellular tail is linked to these pathologies remains unknown.

The 157 amino acid cytoplasmic tail of GlialCAM is dispensable for its trafficking to the plasma membrane and cis-homodimer formation ^1^. While truncation of the cytoplasmic domain did not affect membrane localization, it reduced paracrine adhesion and impaired wound healing, highlighting its role in modulating cell-matrix interactions and cell motility ^1^. Deletion of the GlialCAM cytoplasmic tail disrupts the correct targeting of MLC1, although the localization or activation of the CLC-2 channel is not impacted ^14^.

Here, we have used Cre/lox strategies to generate a characterize a mutant mouse model that expresses GlialCAM containing intact extracellular and transmembrane domains but lacking a cytoplasmic tail. Mutant mice display coordination and motor deficits owing to the development of white matter vacuolation pathologies in the cerebellum. Single cell RNA sequencing and spatial transcript profiling identify differentially expressed genes in glial cells. Furthermore, proteomic experiments reveal that the GlialCAM cytoplasmic tail serves as a hub integrating adhesion complexes with intracellular signaling proteins. This study provides new insight into astrocyte and oligodendrocyte-dependent white matter pathologies and may lead to the identification of potential therapeutic targets to treat brain disorders like MLC.

## RESULTS

### Generation and characterization of the GlialCAM ΔCT mutant mouse model

Analysis of GlialCAM protein expression patterns in the post-natal cerebral cortex reveal enrichment in GFAP^+^ perivascular astrocytes and Olig2^+^ oligodendrocytes (Fig. 1A and B). In the cerebellum we detect GlialCAM expression in GFAP^+^ Bergmann radial glial cells and white matter astrocytes as well as in Olig2^+^ oligodendrocytes and in myelin sheets (Fig. 1A and B). To investigate the function of the GlialCAM cytoplasmic tail in glial cells, we engineered a mouse model that allows for Cre-mediated ablation of the cytoplasmic domain coding sequence without perturbing the extracellular and transmembrane domains (Fig. 1C). The last three exons of the *Hepacam* gene (exons 5-7) encode the 157 amino acid cytoplasmic domain. To inhibit expression of this domain a premature stop codon was introduced at the 3’ end of exon four. To prevent potential dominant-negative effects caused by the engineered stop codon, a cDNA mini-gene comprising the coding sequences for exons four to seven and a polyA tail, flanked by loxP sites, was inserted upstream of exon four. After Flp-mediated removal of the Frt-flanked neomycin selection cassette, the result is a conditional knock-in (ConKI) allele (Fig. 1C). The conditional knockout (ConKO) allele cane be generated through Cre-mediated deletion of the floxed mini-gene sequence, resulting in premature termination of translation at the engineered stop codon. Cre-mediated deletion of the mini-gene was validated in primary brain astrocytes isolated from mouse pups harboring the *Hepacam* ConKI allele. Astrocytes were infected with control adenovirus or adenovirus-Cre and genomic PCR was performed to confirm recombination of the mini-gene sequence in heterozygous (*Hepacam*^ConKI/+^) cells and homozygous (*Hepacam*^ConKI/ConKI^) cells (Fig. 1D).

**Fig. 1.**
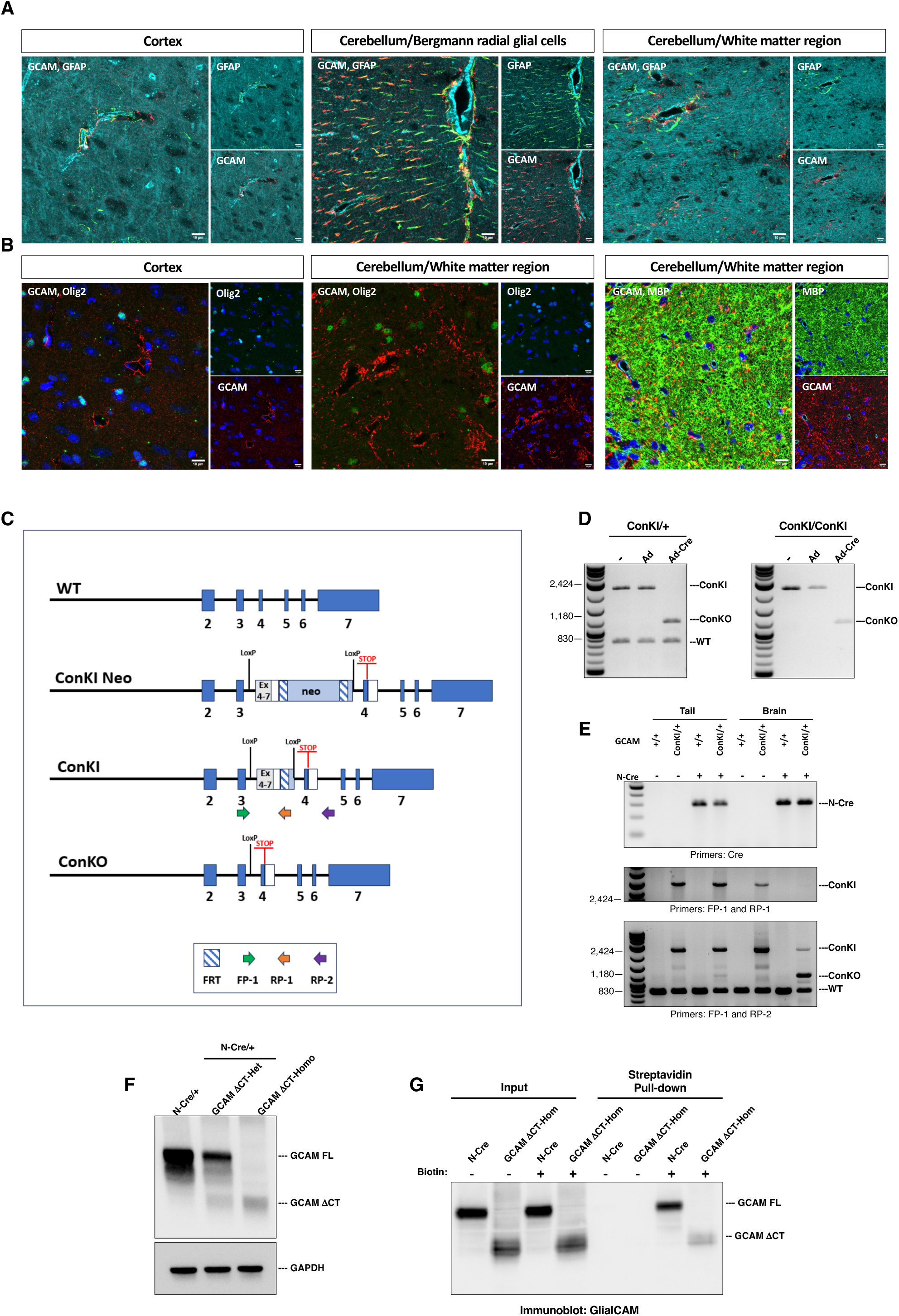
Generation of the GlialCAM ΔCT mutant mouse model. **(A, B);** Representative immunofluorescence images showing GlialCAM (GCAM) expression in GFAP^+^ astrocytes (A) and Olig2^+^/MBP^+^ oligodendrocytes (B) within the cortex and cerebellum of 3.5-month-old mice. Blood vessels are shown in cyan, and nuclei are stained with DAPI. Scale bars, 10 µm. **(C);** Strategy for generating the GlialCAM ΔCT mouse model. A premature stop codon was introduced in exon 4 of the Hepacam gene to prevent expression of the downstream cytoplasmic sequence. To prevent potential dominant negative effects of the engineered stop codon (ConKI-Neo), a cDNA mini-gene comprising coding sequences from exons 4-7 and a polyA tail flanked by loxP sites were inserted upstream of exon 4. The Neo cassette was later removed using Flp recombinase, creating the conditional knock-in allele (ConKI). When Cre recombinase is used to delete the floxed mini-gene sequence, translation is prematurely terminated (due to the STOP codon), resulting in a conditional knockout allele (ConKO) that encodes a truncated protein lacking the cytoplasmic domain. **(D);** PCR analysis of genomic DNA from GlialCAM (GCAM) ConKI-Het (left) or GCAM ConKI-Hom (right) astrocyte cells infected with either control adenovirus (Adeno) or Adeno-Cre. **(E);** PCR analysis of DNA from either tail or brain tissue from P2 GCAM WT (N-Cre^-^:GCAM^+/+^), GCAM ConKI-Het (N-Cre^-^:GCAM^+/ConKI^), N-Cre^+^ (N-Cre^+^:GCAM^+/+^), and GCAM ΔCT-Het (N-Cre^+^:GCAM^+/ConKI^) mice. **(F);** Immunoblot showing GlialCAM (GCAM) full length (FL) and GlialCAM ΔCT expression in brain lysates of control, GCAM ΔCT-Het, and GCAM ΔCT-Homo mice. **(G);** Detergent-soluble lysates from cell surface-biotinylated cortical astrocytes from GCAM ΔCT homozygote pups were incubated with streptavidin-coated beads followed by immunoblotting with antibodies directed against the GlialCAM extracellular domains.

To conditionally delete the *Hepacam* cytoplasmic domain in the brain, *Hepacam*^ConKI/ConKI^ or *Hepacam*^ConKI/+^ female mice were crossed to heterozygous Nestin-Cre/+ male mice ^15^. We have shown previously that Nestin-Cre is active in neuroepithelial cells of the embryonic brain as early as E10-E11 ^16^. The first cross produced offspring with 49.71% GlialCAM mutants, and the second cross yielded 43.48% GlialCAM mutant carriers (30.47% heterozygotes and 26.06% homozygotes). Analysis of Nestin-Cre activities in the post-natal brain using the Rosa26-loxSTOPlox-YFP reporter strain confirmed Cre-dependent YFP expression primarily in GFAP^+^ astrocytes, Olig2^+^ oligodendrocytes, and PDFGRα^+^ OPCs (Supp. Fig. 1). Throughout the study, mice with Nestin-Cre^+^:*Hepacam*^+/+^ (N-Cre) genotype served as controls, Nestin-Cre^+^:*Hepacam*^+/ConKI^ were GlialCAM ΔCT heterozygous; and Nestin-Cre^+^:*Hepacam*^ConKI/ConKI^ were GlialCAM ΔCT homozygous mice. To verify Nestin-Cre-mediated recombination of the ConKI allele, PCR was performed using primers that amplify sequences between the two loxP sites and exon three (Fig. 1E). In GlialCAM ΔCT heterozygous mice, the ConKI band was absent in brain tissue samples, where Cre is active (Fig. 1E). In contrast, this primer set did not reveal ConKI deletion in tail samples due to limited activity of Cre outside of the CNS. To further confirm the generation of the ConKO allele, genotyping was performed using an additional primer set designed to amplify regions flanking the two loxP sites. This primer pair produced a band exclusively in brain samples containing both the ConKI and Nestin-Cre alleles (Fig. 1E).

Immunoblotting of protein extracts from whole brain lysates from four-month-old mice confirmed the expression of the truncated GlialCAM protein in both heterozygous and homozygous GlialCAM ΔCT samples. An antibody against the extracellular region of GlialCAM reveled truncation of GlialCAM protein in heterozygous GlialCAM ΔCT samples and a complete absence of the full-length GlialCAM protein in homozygote GlialCAM ΔCT samples (Fig. 1F). To determine whether C-terminal truncation had any effects on GlialCAM membrane trafficking, cultured neonatal mouse brain astrocytes from GlialCAM ΔCT homozygote mice were surface biotinylated followed by pull-down with streptavidin-conjugated beads and immunoblotting with anti-GlialCAM antibodies against the extracellular region present in both the full-length and truncated proteins (Fig. 1G). We found that GlialCAM ΔCT traffics to the plasma membrane, and it showed slightly reduced plasma membrane levels compared with the GlialCAM full-length (Fig. 1G).

### Mice with C-terminally truncated GlialCAM develop white matter vacuole pathologies

We examined whether GlialCAM truncation leads to any obvious pathologies in vivo. Detailed histopathological analysis revealed vacuole formation in white matter regions of the brain (Fig. 2A-D). Vacuolation was observed in the cerebellar white matter of GlialCAM ΔCT homozygous mice as early as 4 post-natal months (Fig. 2A and C). Vacuoles became progressively more severe with age in the mutant brains, as demonstrated upon comparison of the GlialCAM ΔCT cerebellum at four and 12 months (Fig. 2A and C). Vacuoles were also present in the corpus callosum, although they displayed reduced sizes and were less numerous compared to those in the cerebellar white matter (Fig. 2B and D). While vacuolization in the corpus callosum increased at both four and 12 months of age compared to their respective controls, no obvious differences were observed in the extent of vacuolization between mutants at these two ages. Four-month-old mutants displayed obvious white matter vacuoles within the corpus callosum region like 12- month-old mutants (Fig. 2D). Heterozygous mutant mice did not develop pathological vacuoles in the white matter (data not shown). Therefore, we used GlialCAM ΔCT homozygous mice throughout this study.

**Fig. 2.**
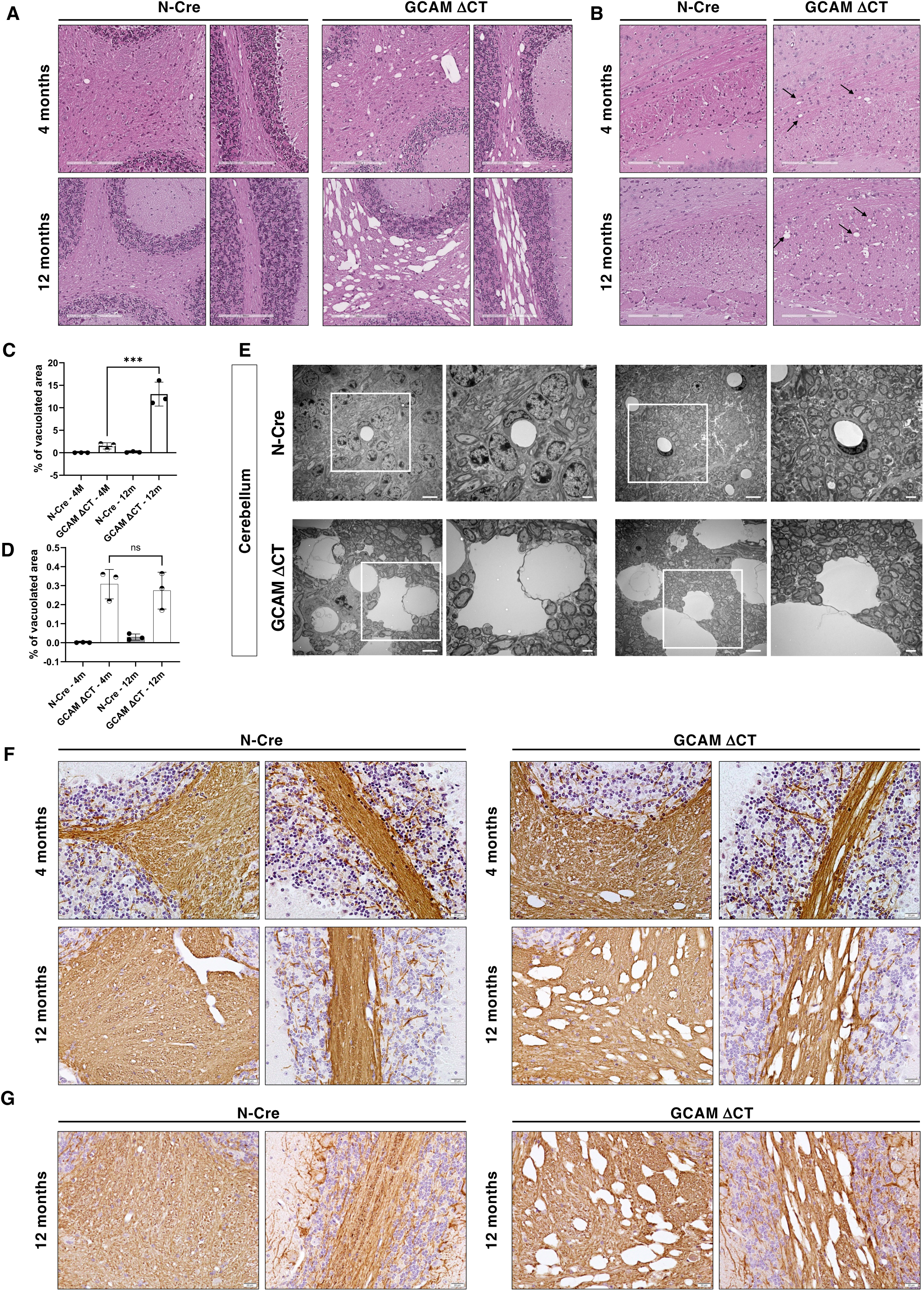
Mice expressing a GlialCAM cytoplasmic domain truncation develop white matter vacuolization pathologies. **(A, B);** H&E-stained sagittal brain sections through the cerebellum (A) and cortex/corpus callosum (B) of four-month-old and 12-month-old controls and GlialCAM (GCAM) ΔCT homozygous mice showing the vacuoles (arrow).Scale bars, 200 µm. **(C, D);** Quantification of the percentage of vacuolated area within the cerebellum (C) and corpus callosum (D) of four month-old and 12-month-old control and GCAM ΔCT mice. Graph bars represent mean ± SD of 3 animals per group. **(E);** Transmission electron microscopy analysis of brain sections from the cerebellum of 12-month-old control and GCAM ΔCT homozygous mice. Boxed areas in the left panels (scale bars, 6 µm) are shown at higher magnification in the right panels (scale bars, 2 µm). **(F);** Anti-MBP immunohistochemical staining of brain sections from the cerebellum of 4-month-old and 12-month-old control and GCAM ΔCT-Homo mice. Scale bars, 20 µm. **(G);** Anti-neurofilament immunohistochemical staining of brain sections from the cerebellum of 12-month-old control and GCAM ΔCT-Homo mice. Scale bars, 20 µm.

Next, ultrastructural analysis of white matter regions in GlialCAM ΔCT and control brains was performed using transmission electron microscopy (Fig. 2E). Vacuoles of variable sizes surrounded by thin myelin sheaths were detected in the GlialCAM ΔCT cerebellar white matter. Immunohistochemistry with anti-MBP and anti-neurofilament antibodies confirmed myelin sheets and neurofilaments surrounding the vacuoles (Fig. 2F and G).

### Behavioral analyses reveal motor coordination and cognitive deficits in GlialCAM ΔCT mice

Given the extensive vacuolization observed in the cerebellum of GlialCAM ΔCT mice, a comprehensive behavioral battery was performed to evaluate the effect of GlialCAM truncation on cognitive function (Novel Object/Place Recognition Test, NOPRT), muscular strength (grip strength), as well as motor coordination and balance (catwalk, beam walk, and rotarod). These assessments were performed on 12-month-old control and GlialCAM ΔCT mice.

The catwalk analysis revealed that the general run parameters in GlialCAM ΔCT and control mice were comparable (data not shown). However, detailed examination of spatial, temporal and interlimb coordination metrics demonstrated significant differences between GlialCAM ΔCT and control mice (Fig. 3A-C). Specifically, among the spatial parameters, GlialCAM ΔCT mice exhibited higher maximum intensity and mean intensity values among the 15 brightest pixels of each paw print in the right front (RF), left front (LF), and one of the front paws (FP). In addition, the mean intensity in the LF and FP paws were elevated in GlialCAM ΔCT mice, while the value for the RF paw remained unchanged (Fig. 3A). Several temporal parameters such as stand times per paw were comparable between two groups, but maximal intensity at paw contact points was significantly greater for RF, LF, and FP paws in mutants, with increased mean intensity at maximum contact of a paw in LF and FP paws (Fig. 3B). Regarding coordination, parameters such as regularity index and phase dispersion showed no differences, but interlimb coupling was significantly diminished in the RF to LH limb of GlialCAM ΔCT mice, whereas coordination across other limbs remained comparable to controls (Fig. 3C).

**Fig. 3.**
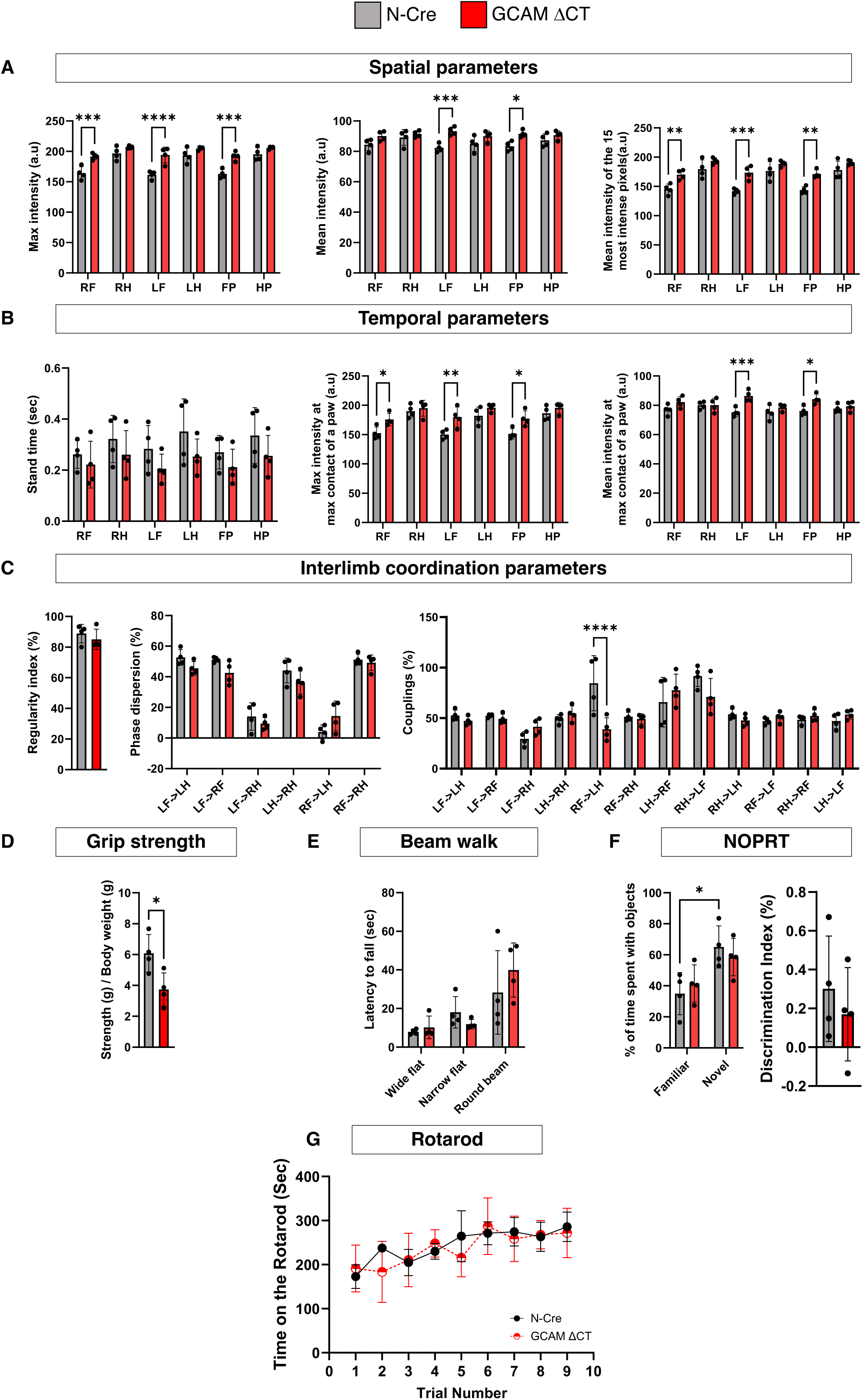
Comprehensive behavioral assessment reveals selective functional impairment in GlialCAM ΔCT mice. **(A-C);** Spatial, temporal and interlimb coordination measurements using the catwalk assay for the GlialCAM ΔCT and control (N-Cre) mice. **(D);** Hindlimb grip strength test of GlialCAM ΔCT compared to control mice. The force that animals used to grip the handle is normalized to their body weight. **(E);** The beam walking test reveals lack of statistically significant differences in balance between control and mutant mice. **(F);** The NOPRT assay reveals impaired novelty recognition in GlialCAM ΔCT mice, as well as no significantly robust group differences. **(G);** Rotarod analyses of 12-month-old control and GCAM ΔCT-Homo mice (n=3). The time that mice stayed on the rod is plotted against the trial number. All data were analyzed using unpaired t test or two-way ANOVA followed by Bonferroni multiple comparisons. All data are presented as mean ± SD, n=4. Right front paw (RF), right hind paw (RH), left front paw (LF), left hind paw (LH), one of the front paws (FP), and one of the hind paws (HP).

While GlialCAM ΔCT mice exhibited less hindlimb grip strength compared with controls (p value=0.0276) (Fig. 3D), no differences were found between GlialCAM ΔCT mice and control in the beam walk test (Fig. 3E). In the NOPRT assay, control mice spent significantly more time exploring the novel object compared to the familiar one (adjusted p-value=0.0269), indicating intact recognition memory. In contrast, GlialCAM ΔCT mice did not show such a preference (adjusted p-value =0.2878), revealing impaired novelty recognition (Fig. 3F). Consistent with this, the discrimination index (DI) value trended higher in control mice (mean DI = 0.3012 ± 0.2714 SD) compared to GlialCAM ΔCT mice (mean DI = 0.17 ± 0.2401 SD), but these values were not significantly different from zero (WT: t(3)=2.22, P Value=0.113; Mut: t(3)=1.416, P Value=0.2518), nor did they differ significantly between groups (Fig. 3F). Thus, while % exploration time indicates intact recognition memory in control mice and impaired recognition in GlialCAM ΔCT mice, the DI analysis did not reveal statistically robust group differences. Furthermore, no significant differences in rotarod performance were observed between GlialCAM ΔCT mice and controls (Fig. 3G).

### GlialCAM cytoplasmic domain truncation does not abrogate BBB integrity

GlialCAM is expressed in perivascular astrocytes (Fig. 1A), which play key roles in regulating BBB maturation and homeostasis; therefore, we examined GlialCAM ΔCT mice for evidence of defects in endothelial barrier integrity. Cadaverine-FITC is a 640 Dalton tracer that does not normally enter the brain parenchyma unless the BBB is compromised ^17^. Indeed, using the middle cerebral artery occlusion (MCAO) model of ischemic stroke, which impairs endothelial barrier integrity, we detected robust extravasation of cadaverine (Supp. Fig. 2A). However, when we compared 12 month-old GlialCAM ΔCT and control mice, no differences in cadaverine leakage into the brain parenchyma were detected (Supp. Fig. 2B and C). Additionally, transmission electron microscopy analysis revealed no obvious cytoarchitectural abnormalities in the blood vessels of GlialCAM ΔCT mice (Supp. Fig. 2D).

### Profiling gene expression changes following C-terminal truncation of GlialCAM

Various transmembrane cell-cell adhesion proteins, like cadherins, can regulate signaling events leading to alterations in gene expression ^18^. To determine if GlialCAM has roles in regulating transcriptional responses in the brain, we performed single cell whole transcriptome profiling of the cerebral cortex and cerebellum from 12 month-old mice. A total of 22,969 cells were sequenced in the cerebral cortex sample. This included 12,153 cells from wild-type mice (n=2) and 10,816 cells from GlialCAM ΔCT mice (n=2). Uniform manifold approximation and projection (UMAP) was employed to visualize the distribution of each cell type (Fig. 4A). We used known cell type-enriched genes to determine the identities of the clusters generated by UMAP (Fig. 4B). As shown in Fig. 4C, *Hepacam* transcripts were enriched in astrocytes (clusters 1, 13, and 21), oligodendrocytes (clusters 0 and 6), and OPCs (cluster 14).

**Fig. 4.**
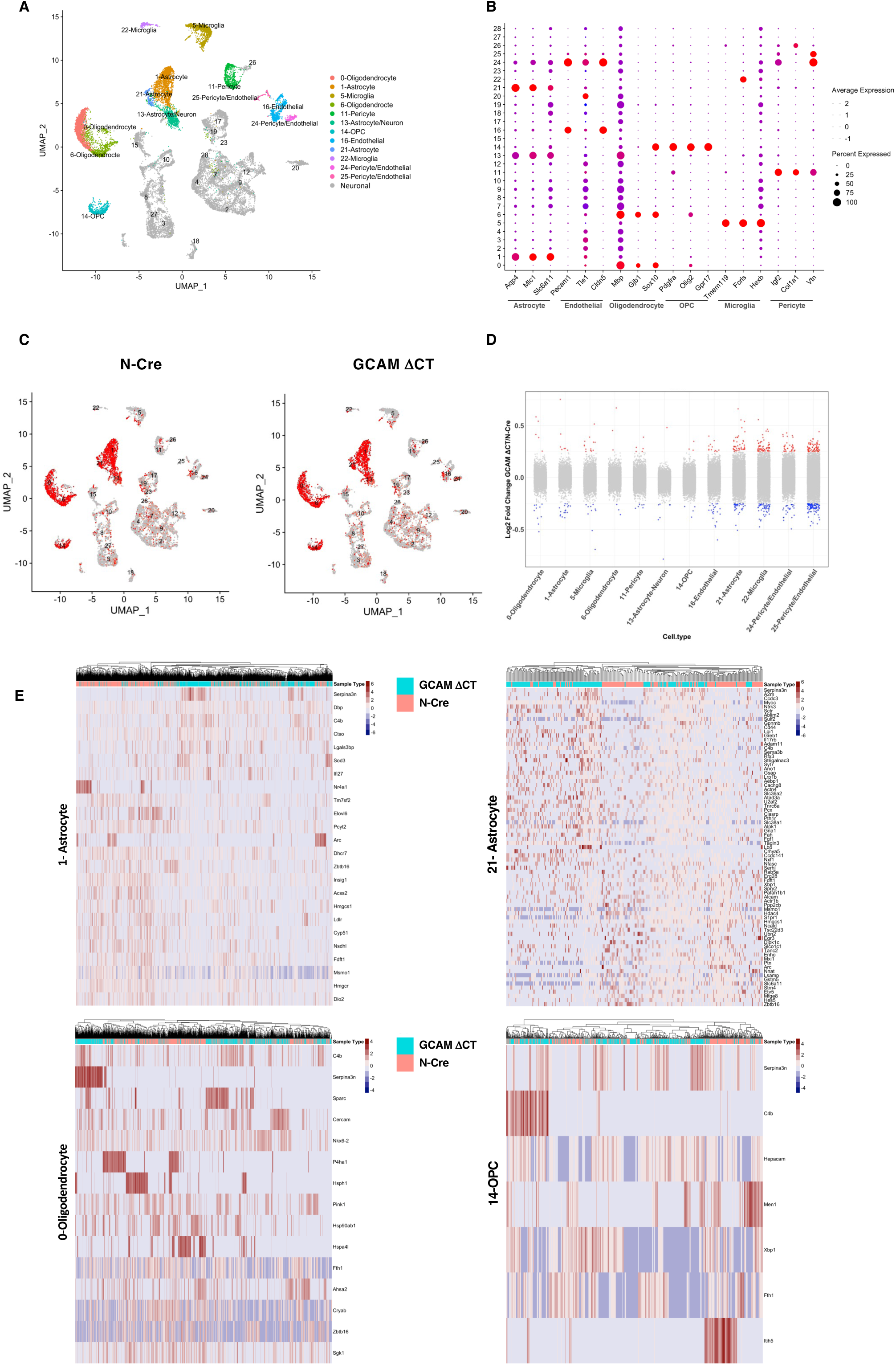
Analysis of gene expression changes associated with GlialCAM C-terminal truncation in cerebral cortex cells from scRNA-seq data. **(A);** UMAP visualization of the 29 clusters identified in the control and GlialCAM (GCAM) ΔCT cortical cells. **(B);** Dot plot showing the distribution of expression levels of well-known representative cell-type-enriched marker genes across all clusters. **(C);** UMAP visualizing the expression distribution of Hepacam across different cell-types in control and GCAM ΔCT cortical cells. **(D);** Strip chart illustrating the logarithmic fold changes (log FC) of all detected genes across cell types in GCAM ΔCT samples relative to controls. Genes that are significantly up-regulated and down-regulated (P value < 0.05 and 0.25 ≤ FC ≤ −0.25) are color coded for each cluster. **(E);** Heatmap visualizing log-normalized, scaled expression of differentially expressed genes (DEGs) identified within *Hepacam*-enriched clusters (astrocyte, oligodendrocyte and OPC_)._

To investigate the molecular genetic effects of GlialCAM cytoplasmic domain truncation, we next performed differential gene expression analysis. Across the dataset, 1962 genes were detected, of which 385 genes showed significant expression changes linked to GlialCAM ΔCT in at least one cell type (adjusted p-value threshold of ≤ 0.05), with an average log-fold difference of at least 0.25 between the two groups. (Fig. 4D). More than half of the differentially expressed genes (DEGs) were shared across various cell types, even in those that lacked Hepacam transcripts, suggesting that the GlialCAM cytoplasmic domain controls paracrine signaling responses. In addition to these common gene expression changes, GlialCAM ΔCT also induced cell type-specific responses. Among the non-neuronal cell populations analyzed, microglia cells exhibited the greatest number of unique DEGs, accounting for 73.9% of the total DEGs within that cluster. This was followed by the astrocyte population (cluster 1), which contributed 70.8% of the unique DEGs (Fig. 4D).

To better understand the effects of gene expression variation linked to the cytoplasmic domain we focused on DEGs specific to cell types that express endogenous Hepacam. Among these, Cercam was unique to only oligodendrocyte clusters, where its expression was upregulated by 1.2-fold in cells harboring GlialCAM ΔCT. In the brain Cercam is predominantly expressed in oligodendrocytes, although its precise biological functions are unclear ^19^. Additionally, genes associated with the inflammatory response, including complement factor C4b and the serine peptidase inhibitor Serpina3n, were upregulated in all *Hepacam*-enriched cell clusters. Serpina3n expression was also upregulated in several neuronal clusters that did not express *Hepacam*, suggesting a non-cell autonomous involvement in the neuroinflammatory process. In contrast, the elevated expression of C4b was restricted to *Hepacam*-enriched clusters, indicating a more cell type-autonomous role in the inflammatory response within these populations (Fig. 4E).

GO annotation analysis indicated that astrocyte cluster 1 transcriptional changes are linked to cholesterol biosynthesis and lipid metabolism pathways, crucial for maintaining membrane structure and cell signaling. Several key genes involved in these pathways, such as Hmgcr, Msmo1, Fdft1, Nsdhl, Cyp51, Ldlr, Hmgcs1, Acss2, Insig1, Dhcr7, Pcyt2, Elovl6, and Tm7sf2, were downregulated in astrocyte cluster 1 from GlialCAM ΔCT mice (Fig. 4E). In astrocyte cluster 21, genes involved in innate immunity and inflammation (C4b, A2m, and Serpina3n), as well as genes involved in ECM and cytoskeleton organization (Sulf2, and Myoc) were upregulated in GlialCAM ΔCT astrocytes (Fig. 4E). DEGs within the OPC subpopulation were linked to pathways essential for cellular stress responses (Fth1, and Xbp1), ECM stabilization and cell adhesion (Itih5, and Hepacam), as well as cell cycle regulation (Men1) (Fig. 4E). In oligodendrocyte cluster 0, the gene encoding cysteine-rich acidic matrix-associated protein (Sparc) which is involved in ECM remodeling along with the ones essential for myelin formation and cell adhesion (Nkx6-2 and Cercam) were upregulated in GlialCAM ΔCT astrocytes. The downregulated genes were mostly enriched in stress-related responses and protein folding processes, such as Sgk1, Cryab, Ahsa2, Hspa4l, Hsp90ab1, Pink1, and Hsph1 (Fig. 4E).

DEGs in the cerebellum were analyzed in a total of 35,599 single cells derived from 12- month-old wild-type mice (n=2) (17,370 cells) and age-matched GlialCAM ΔCT mice (n=2) (18,229 cells). Using known cell type markers, we annotated 21 distinct clusters identified through UMAP analysis (Fig. 5A and B). *Hepacam* expression was specifically enriched in astrocytes (clusters 6, and 8) and oligodendrocytes (cluster 5) (Fig. 5C). DEG analysis detected 1505 genes, of which 325 genes exhibited statistically significant alterations in expression linked to GlialCAM truncation, with an average log-fold difference of at least 0.25 between the two groups (Fig. 5D). The majority of DEGs in cerebellar cells did not overlap with other cell clusters. Among the *Hepacam* enriched clusters, astrocyte cluster 6 subpopulation exhibited the highest number of DEGs, with 83.72% of them being unique to this cluster. Among the downregulated DEGs there were ones involved in metabolism pathways (Etnppl, Txnip, Aass, and Acsl3) and ECM organization and cell adhesion (Pxdn). Upregulated DEGs in GlialCAM ΔCT cells had links to cytoskeletal dynamics (Flna, Kif21b, and Cnn2) and ECM organization (Fmod, Fbln5, and Adamts1). While some of the upregulated genes in astrocyte cluster 8 from GlialCAM ΔCT mice were shared with cluster 6 astrocytes and were also linked to ECM organization (Fmod, and Adamts1), Plcb1 expression was uniquely elevated in this cluster (Fig. 5E). Additionally, we found that downregulated DEGs in GlialCAM ΔCT oligodendrocytes were enriched in cytoskeletal dynamics and cell motility (Stmn1, and Tmsb4x), metabolism pathways (Glul, Fth1, and Adi1), and stress responses (Cryab, Hsph1, Hspa8, Hsp90aa1, and Hsp90ab1); and upregulated genes were enriched in the immune response (C4b, and Serpina3n) and ECM remodeling (Sparc) (Fig. 5E).

**Fig. 5.**
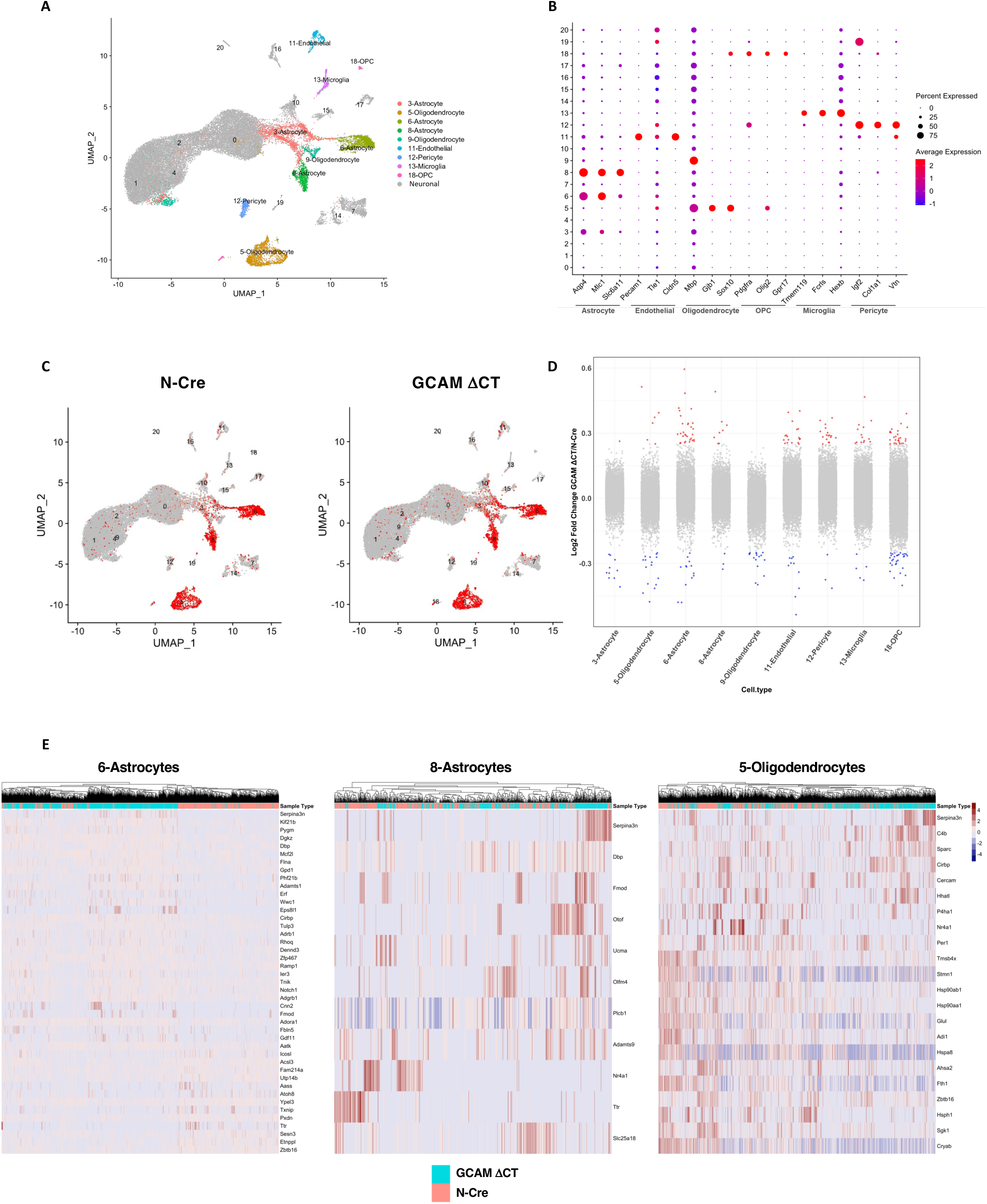
Analysis of gene expression changes associated with GlialCAM C-terminal truncation in cerebellum cells from scRNA-seq data. **(A);** UMAP visualization of the 21 clusters identified in the control and GlialCAM (GCAM) ΔCT cerebellum cells. **(B);** Dot plot showing the distribution of expression levels of well-known representative cell-type-enriched marker genes across all clusters. **(C);** UMAP visualizing the expression distribution of Hepacam across different cell-types in control and GCAM ΔCT cerebellar samples. **(D);** Strip chart illustrating the logarithmic fold changes (log FC) of all detected genes across cell types in GCAM ΔCT cortical cells relative to controls. Genes that are significantly up-regulated and down-regulated (P value < 0.05 and 0.25 ≤ FC ≤ −0.25) are color coded for each cluster. **(E);** Heatmap visualizing log-normalized, scaled expression of differentially expressed genes (DEGs) identified within *Hepacam*-enriched clusters (astrocyte, and oligodendrocyte_)._

Levels of Serpina3n were elevated in cerebellar cells from GlialCAM ΔCT mice, with expression restricted to Hepacam-enriched cells, specifically astrocytes and oligodendrocytes. Immunofluorescence staining confirmed increased Serpina3n protein in both cortex and cerebellum regions of GlialCAM ΔCT mice compared to control mice (Fig. 6). GlialCAM ΔCT mice exhibited significantly higher Serpina3n expression in GFAP^+^ astrocytes (Fig. 6A and B), Olig2^+^ oligodendrocytes (Fig. 6C and D) and PDFGRα^+^ OPCs (Fig. 6E), corroborating our scRNA-seq findings of Serpina3n upregulation within these glial subtypes.

**Fig. 6.**
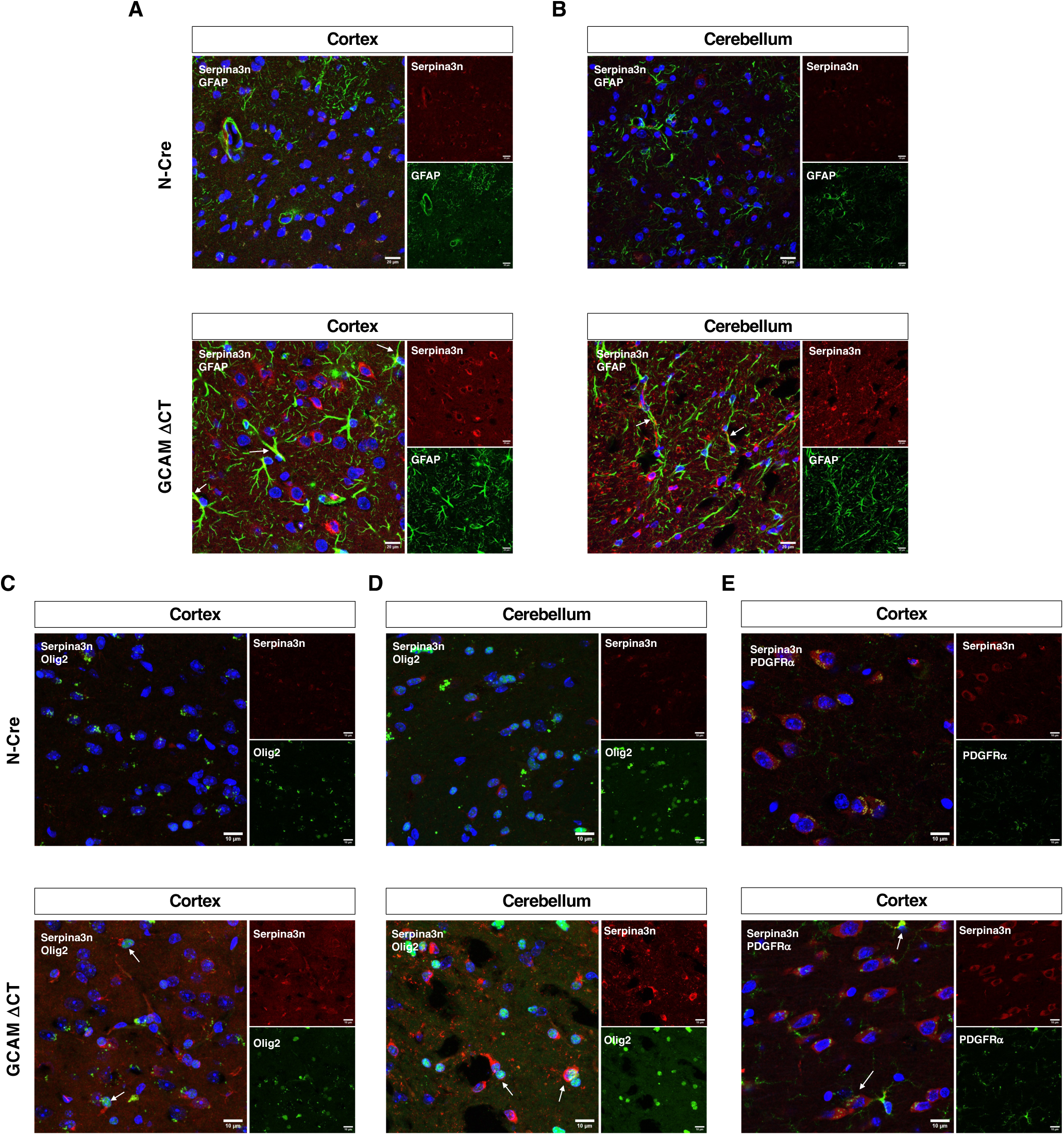
Serpina3n protein is upregulated in GFAP^+^, Olig2^+^, and PDGFRα^+^ glial cells in GlialCAM ΔCT mice. **(A-E);** Immunofluorescence staining was performed on sagittal sections through the cortex and cerebellum from 12-month-old control and GlialCAM (GCAM) ΔCT mice. Sections were stained with anti-Serpina3n antibodies (red) together with anti-GFAP antibodies (A, B),, or anti-Olig2 antibodies (C, D). Additional Immunofluorescence staining with anti PDFGRα and Serpina3n was performed on cortical sections (E). Note the increased expression of Serpina3n protein in glial cells of the mutant brain. Cells co-expressing Serpina3n and indicated cell markers are shown by arrow.

Next, in situ single cell spatial transcriptomics was employed to examine the distribution of different cell types across cortical and cerebellum regions in GlialCAM ΔCT and control samples. The Xenium Mouse Brain Gene Expression panel was originally designed to target 247 genes, covering markers for seven primary cell types: astrocytes, oligodendrocytes, endothelial cells, pericytes, macroglia, fibroblast, and neurons. To enhance its utility, the panel was customized with the addition of 49 genes, primarily associated with activated astrocytes, expanding it to a total of 296 genes (Supp. Table1).

Using UMAP and known cell-specific markers, we identified 20 distinct clusters across cortical regions (Supp. Fig. 3A and B). Analysis of non-neuronal cell types revealed only minor differences in their relative abundance. The fold change in cell type proportion in GlialCAM ΔCT mouse compared with the control was 0.98 in cluster 4-astrocyte, 0.84 in cluster 0-oligodendrocyte, 0.8 in cluster 10-oligodendrocyte, 0.9 in cluster 17-fibroblast, 1.21 in cluster 5-endothelial, and 1.36 in cluster 8-microglia (Supp. Fig. 3C). We initially focused on *Hepacam*-expressing cell types identified through our scRNA-seq data. The analysis revealed that the spatial distribution of astrocytes was comparable between GlialCAM ΔCT and control samples across different cortical regions (Supp. Fig. 3D). However, a distinct cellular pattern was observed for oligodendrocyte clusters between GlialCAM ΔCT and control samples. UMAP analysis identified three oligodendrocyte clusters, each exhibiting a unique distribution across the cortical region. In both GlialCAM ΔCT and control samples, cluster 11 oligodendrocytes were found in both the cortical regions and the corpus callosum, cluster 0 oligodendrocytes were mostly confined to the corpus callosum, and cluster 10 oligodendrocyte cells were restricted to the cortex (Supp. Fig. 3E). All three oligodendrocytes cells displayed a distinct spatial organization in the GlialCAM ΔCT sample. While the relative abundance of the 11-oligodendrocyte cluster was similar between the mutant and control, these cells exhibited higher cell density in the mutant frontal cortex in comparison with the control. While the relative abundance of both 0-oligodendrocyte and 10-oligodendrocyte clusters were lower in the GlialCAM ΔCT compared to the ones in the control sample, these oligodendrocyte clusters exhibited lower cell density in the frontal region of the GlialCAM ΔCT cortex compared to the control sample (Supp. Fig. 3E).

In addition, we observed differences in neuronal patterning between GlialCAM ΔCT and control samples across various layers in the frontal cortex. In the GlialCAM ΔCT mouse, the neuronal cell population in layer V (clusters 7 and 12) was more widely distributed compared to the corresponding layer in the control sample (Supp. Fig. 3F). Similarly, layer V/VI (cluster 6) exhibited higher cell density in the GlialCAM ΔCT frontal cortex despite having a comparable relative abundance in the cortical region between the GlialCAM ΔCT and control samples (Supp. Fig. 3G). Conversely, layer VI (cluster 2) in the GlialCAM ΔCT frontal cortex exhibited reduced cell density (Supp. Fig. 3G). Furthermore, in the GlialCAM ΔCT frontal cortex, the boundary between layer V (cluster 7 and 12) and VI (cluster 2) was unclear (Supp. Fig. 3H). In addition, neurons in layer 6b (cluster 18) were fewer in GlialCAM ΔCT cortex in comparison to the control sample, and this difference was more significant in the frontal cortex (Supp. Fig. 3 D-H). To more clearly illustrate cell distribution differences between the control and GlialCAM ΔCT frontal cortex, spatial density maps for each cluster are provided in Supp. Fig. 4 A-I.

We also performed spatial transcriptomic analysis to assess the distribution of various cell types in cerebellar regions of GlialCAM ΔCT and control mice. Using cell type markers, we identified 18 distinct cell clusters (Fig. 7A, and Supp. Fig. 5A) with no significant GlialCAM-dependent differences in cell proportions (Fig. 7B). Focusing on non-neuronal cells, we observed no significant differences in the overall proportions of various cell types. However, a slight increase in fibroblasts localized at the intersection of cerebellar folia was noted in the GlialCAM ΔCT mouse compared to the control (Supp. Fig. 5B). This difference was more pronounced in the white matter, where the proportion of fibroblasts in the GlialCAM ΔCT mouse was more than two-fold higher than in the control (10% vs 4.5%). Furthermore, microglia cell numbers were elevated in the white matter of the GlialCAM ΔCT cerebellum compared to the control (Supp. Fig. 5C). This increase in microglia was confirmed by immunofluorescence staining of cerebellar regions with anti-Iba1 antibody (Fig. 8A). Iba1^+^ cells were comparable in GlialCAM ΔCT and control cortex regions, consistent with our scRNAseq data (Fig. 8B).

**Fig. 7.**
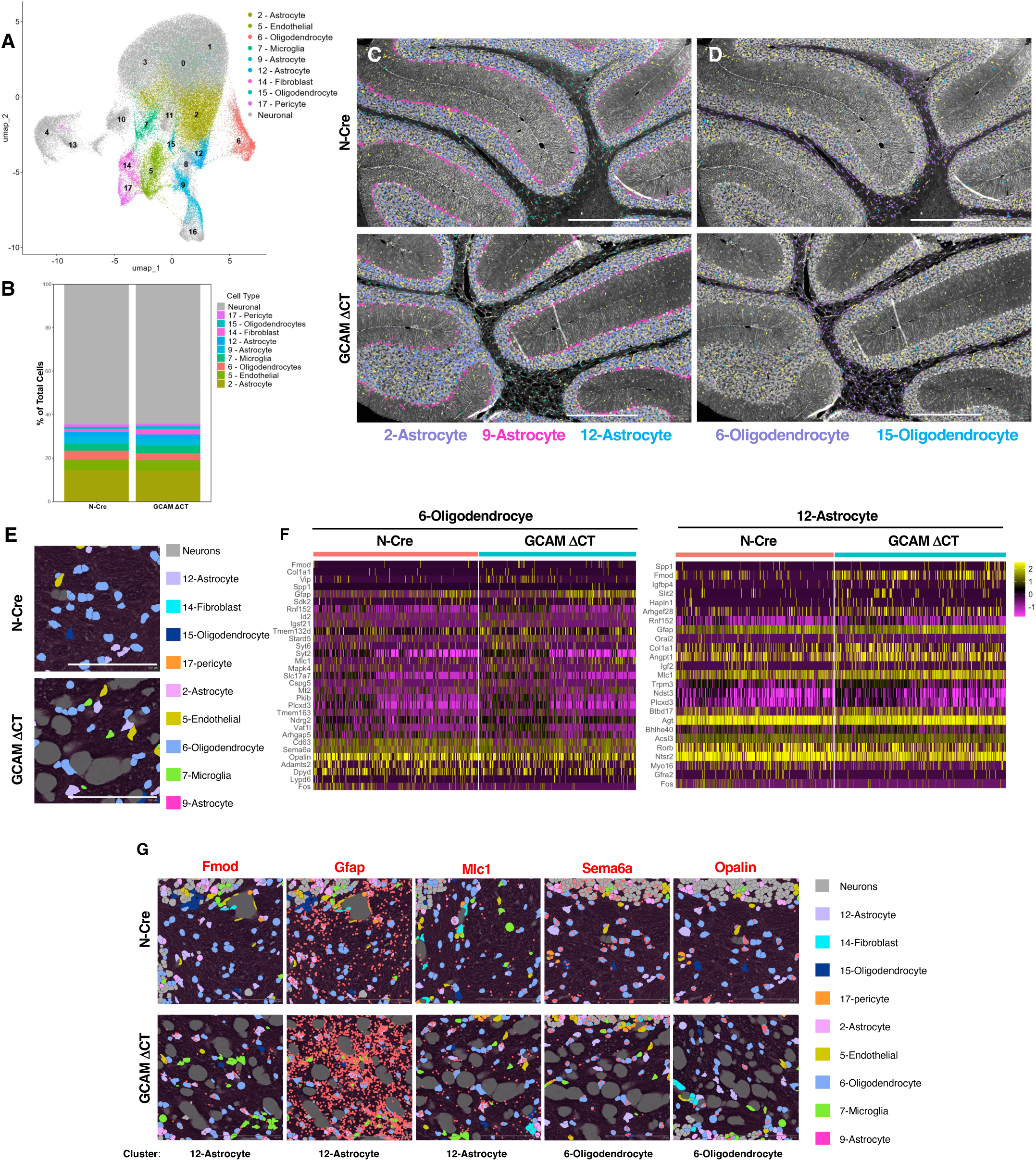
In situ spatial transcript analysis of cerebellar regions in GlialCAM ΔCT and control mouse brains. **(A);** UMAP visualization of 18 clusters identified by Xenium analysis in the cerebellum region of control and GlialCAM (GCAM) ΔCT mice. **(B);** Bar plot showing the proportion of each cell type across all clusters identified within the cerebellum of control and GCAM ΔCT mice. **(C, D);** Special transcriptomic data from 12-month-old control and GCAM ΔCT mouse cerebellum illustrating the spatial distribution of cells within astrocyte clusters (2, 9, and 12) **(C)** and oligodendrocyte clusters (6 and 15) **(D)**, as well as the endothelial cluster (yellow), scale bar=500um. **(E);** Higher magnification view of a representative white matter region illustrating spatial arrangement of distinct cell types, scale bar=100um. **(F);** DEGs identified in cluster 12 astrocyte and cluster 6 oligodendrocyte. **(G);** Spatial *in situ* transcript localization demonstrating expression patterns of Fmod, Gfap, Mlc1, Sema6a, and Opalin genes.

**Figure 8.**
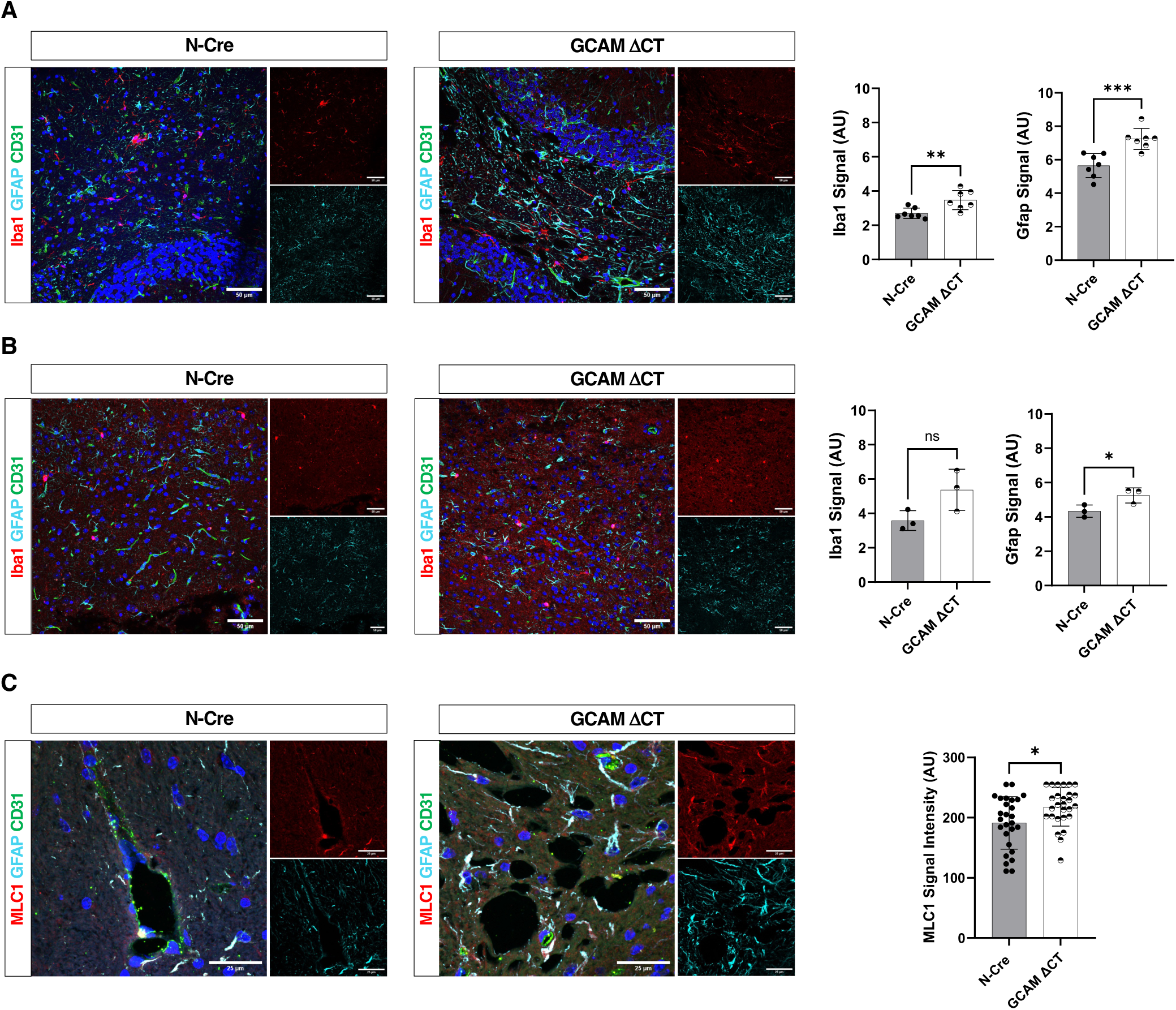
GlialCAM ΔCT mice displayed pronounced astrogliosis with upregulated MLC1 expression in astrocytes and increased Iba-1 immunoreactivity in the cortex and cerebellar white matter. **(A-C);** Immunofluorescence staining was performed on cerebellum (A, C) and cortex (B) sections from 12-month-old control and GlialCAM (GCAM) ΔCT mice. Sections were stained with anti-CD31 (green), anti-Iba-1 (red), and anti-GFAP (cyan) (A and B). Additional staining was performed with anti-CD31 (green), anti-MLC1 (red), and anti-GFAP (cyan) on cerebellum sections (C). Nuclei are counterstained with DAPI. Scale bars, 50 and 25 μM.

We identified three distinct astrocyte clusters in the cerebellum, each showing a different spatial distribution. In both GlialCAM ΔCT and control samples, cluster 2 astrocytes were restricted to the internal granule cell layer. Cluster 9 astrocytes corresponded to Bergmann radial glia, and cluster 12 astrocytes were found within both the internal granule cell layer and cerebellar white matter (Fig. 7C). We also identified two different oligodendrocyte clusters. Cluster 6 oligodendrocytes were mostly restricted to cerebellar white matter regions and cluster 15 oligodendrocytes were mostly located within the internal granule cell layer (Fig. 7D). The spatial distribution of *Hepacam*-enriched clusters, however, were comparable between GlialCAM ΔCT and control brains. Analysis of the mutant white matter regions, which contained vacuoles, revealed two prominent glial populations: cluster 12 astrocytes and cluster 6 oligodendrocytes (Fig. 7E). Both clusters exhibited DEGs associated with ECM remodeling. Notably, Spp1, Fmod, Col1a1, and Hapln1 were upregulated in cerebellar astrocytes in GlialCAM ΔCT mice compared to controls (Fig. 7F). Similarly, Fmod, Col1a1, Adamts2, and Sema6a were enriched in cerebellar oligodendrocytes of GlialCAM ΔCT mice (Fig. 7F and G). Consistent with our immunofluorescence data (Fig. 8A and B), Gfap, a marker for astrocytes was elevated in GlialCAM ΔCT mice cortical and cerebellar regions. Additionally, Oplain, which is involved in oligodendrocyte differentiation and myelination, was upregulated in cluster 6 oligodendrocytes in the GlialCAM ΔCT cerebellum (Fig. 7F, and G). The GlialCAM-interacting partner Mlc1 was also upregulated in cerebellar white matter astrocytes of GlialCAM ΔCT mice (Fig. 7F, and G). This finding was further evaluated by double immunofluorescence labeling for GFAP and MLC1 in white matter areas (Fig. 8C). In GlialCAM ΔCT mice, Mlc1 protein levels were elevated in GFAP^+^ cells relative to controls. MLC1 was also present in astrocyte processes surrounding vacuoles within mutant white matter, although its distribution in astrocytes was similar between GlialCAM ΔCT and control groups (Fig. 8C). Mlc1 upregulation may reflect the reactive state of the astrocytes, since we have shown higher levels of Mlc1 in astrocytes in response to brain trauma ^20^. Collectively, these findings highlight a role for the C-terminal tail of GlialCAM in regulating ECM-associated signaling pathways, underscoring its importance in maintaining white matter structural integrity within the cerebellum.

### Proteomics analysis identifies GlialCAM cytosolic tail signaling partners

To further examine downstream signaling networks associated with the cytosolic tail of GlialCAM, we performed antibody-based reverse-phase protein array (RPPA) profiling on astrocytes isolated from neonatal cortices. Of the 499 phospho-specific and total protein targets analyzed, 35 showed significant expression changes in the absence of the GlialCAM cytosolic tail (q value <0.01; Supp. Fig. 6A and B). After applying a log2 fold change filter (0.5 < log_2_FC <-0.5), 7 proteins remained (Supp. Fig. 6A) and mapped to pathways involved in mitochondrial biogenesis/dynamics (TFAM, MFN1), DNA damage/metabolic processes (RPA2, LIG4), and RTK/Src-regulated ECM remodeling (PTPN12, MMP14) ^21,22^, with SOX2 indicating transcriptional reprogramming toward a reactive state ^23^. RPPA findings for PDK1, CSK, MMP14, and SOX2 were validated by western blotting (Supp. Fig. 6C and D).

To identify GlialCAM cytosolic binding partners that mediate adhesion and signaling functions, we performed GST pull-down assays using a GST-tagged mouse GlialCAM cytosolic tail (GC-Cyto) and control GST (EV), followed by mass spectrometry. GST-fusion proteins were incubated with mouse cerebellum lysates, while buffer-only incubations served as additional controls (Fig. 9A). To maximize protein identification sensitivity, gel lanes were fractionated around 50 kDa and processed separately. Comparative analysis of protein abundances between GC-Cyto and EV, as well as GC-Cyto and buffer only controls, revealed 59 proteins present in both fractions, 169 unique to the >50 kDa fraction (Up), and 77 unique to the <50 kDa fraction (Down) (Fig. 9B) (Supp Table.2). When visualized, these datasets distributed across four quadrants, reflecting the degree of enrichment in GC-Cyto relative to control conditions (Fig. 9C).

**Fig. 9.**
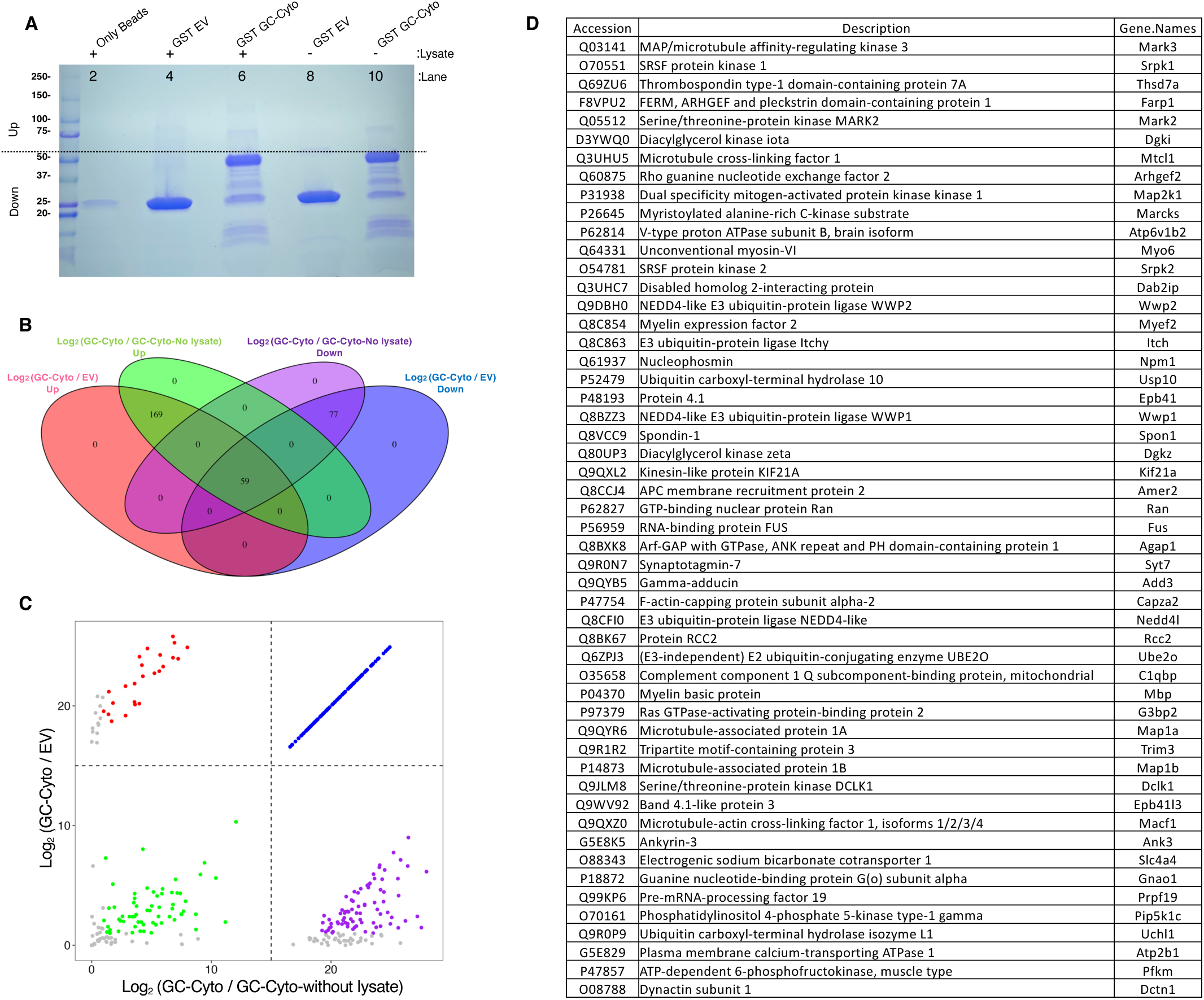
Mass spectrometry based proteomic profiling of GlialCAM ΔCT and control cerebellum samples. **(A)**; Coomassie-stained protein gel of immunoprecipitated samples in the presence or absence of mouse cerebellum lysate. **(B)**; Venn diagrams showing overlap of protein hits across four fractions separated at 50 kDa. **(C);** Scatterplot illustrating relative protein abundance in GC-Cyto versus EV or GC-Cyto-no lysate samples. Protein with log2 fold change of less than 2 are shown in gray. **(D);** Representative list of identified GlialCAM cytosolic interacting partners.

A total of 111 proteins were identified in the <u>top right</u> quadrant, representing proteins highly enriched in the GC-Cyto sample compared with both controls. Several of these hits are associated with actin cytoskeleton organization and dynamics (Thsd7a, MTCL1, MARCKS, FARP1, MYO6), kinase signaling cascades (MARK2/3, DGKI, MAP2K1, SRPK1/2), the subunit of V-type proton ATPase, Atp6v1b2, the Rho guanine nucleotide exchange factor 2, ARHGEF2, and the previously identified GlialCAM interactor the synaptosome-associated protein 25 ^11^. MAP2K1 is a key upstream activator of ERK1/2. In oligodendrocytes Map2k1 is known to promote myelination ^24^.

The stability and dynamics of compact myelin are partly regulated through MAPK/ERK-mediated MBP phosphorylation. Phosphorylation of MBP inhibits its ability to promote cytoskeleton components polymerization and membrane adhesion ^25^.

A total of 36 proteins were identified in the <u>top left</u> quadrant, representing proteins highly enriched in GC-Cyto compared with EV. Among these, 24 proteins were significantly enriched in GC-Cyto relative to GC-Cyto without lysate (log₂FC > 1). While most hits were associated with transcription, translation, and RNA processing, the scaffold protein DAB2 interacting protein, known to regulate AKT1 and MAP3K5 signaling, was also identified as a GC-Cyto-interacting partner.

A total of 117 proteins were identified in the <u>bottom right</u> quadrant, representing proteins highly enriched in GC-Cyto compared with GC-Cyto without lysate. Of these, 79 proteins were significantly enriched in GC-Cyto versus EV (log2FC>2). Among the hits, C1QBP (p32), which acts upstream of complement of C4-B, was notable as loss of *C1qbp* in the CNS has been associated with leukoencephalopathy characterized with white matter vacuolization and oligodendrocyte loss ^26^. Several components of the ubiquitin system were detected including the E3 ubiquitin protein ligase, WWP2, ITCH and NED4L; the deubiquitinating enzyme USP10; and the E2/E3 hybrid ubiquitin-protein ligase, UBE2O. Additional hits included transcriptional repressor of MBP, MYEF2, and the NPM1, which regulates cellular volume by modulating protein synthesis ^27^. Multiple small GTPase signaling proteins and cytoskeleton modulators were identified, such as AGAP1, the RasGRP inhibitor DGKZ ^28^, along with EPB4L, KIF21A, RAN, ADD3, CAPZA2, and RCC2. Signaling scaffolds and pathway modulators were also identified such as AMER2, a negative regulator of Wnt signaling. Additional notable hits comprised the ECM/Cell adhesion protein, SPON1; the RNA-binding protein FUS, whose absence in oligodendrocytes has been linked to increased myelin deposition and cholesterol and AKT activation ^29^; the vesicle trafficking Ca^2+^ sensor SYT7; and the serine/threonine protein kinase DCLK1.

A total of 295 proteins were identified in the <u>bottom-left</u> quadrant of the plot. Among these, 65 proteins were significantly enriched in both GC-Cyto versus EV and GC-Cyto versus GC-Cyto without lysate (log₂FC > 2). These included myelin structural components such as MBP, and the Ras GTPase-activating protein-binding protein G3BP2. Several cytoskeletal regulators were also enriched, including MAP1A/B, EPB41L3, MACF1, and DCTN1. Ion transport and homeostasis– related proteins were prominent, comprising the Kv1.1 channel regulator ANK3, the sodium– bicarbonate cotransporter Slc4a4, the GPCR downstream effector GNAO1, and the plasma membrane Ca²⁺ transporter ATP2B1. The dataset also included signaling enzymes such as the phosphatidylinositol-4-phosphate 5-kinase type-1 gamma (Pip5k1c) and the glycolytic enzyme phosphofructokinase muscle type PFKM. Among these hits, Slc4a4 and MPB have previously identified as GlialCAM interacting partners ^11^ (Fig. 9D).

## DISCUSSION

This study is the first to address functions for the GlialCAM cytoplasmic domain in vivo using a conditional knockout mouse model expressing a C-terminal truncated form of GlialCAM in brain glial cells. GlialCAM C-terminal truncation leads to white matter vacuolization, a phenotype like that observed in MLC patients. Vacuoles were detected mainly within the white matter regions of the GlialCAM ΔCT cerebellum and, to a lesser extent in the corpus collosum. Integrity of the BBB was not compromised following cytoplasmic domain truncation. The GlialCAM cytosolic tail truncation resulted in widespread astrogliosis, evident by increased GFAP expression. There was upregulated expression of the microglia marker Iba1 in the cerebellum. Single cell transcriptome analysis revealed that components of ECM remodeling were among the most significantly affected pathways due to the cytoplasmic truncation.

In astrocytes GlialCAM can function as a chaperone, mediating the localization of MLC1 and ClC-2 at the plasma membrane ^3,8^. Loss of Mlc1 disrupts the localization of both GlialCAM and ClC-2, highlighting the functional interdependency among these interacting partners ^9^. These findings suggest that the loss of ClC-2 serves as the primary cause of the shared pathologies observed in the GlialCAM and Mlc1 mutant models. Cerebellar white matter vacuolization and edema have been reported in null mouse models for Hepacam, Mlc1, and Clc-2. Deletion of Hepacam in Clc-2 null mice exacerbates vacuolization in Clcn2^-/-^ mice, indicating that GlialCAM may play distinct and independent roles in the formation of vacuoles ^9^. Additionally, GlialCAM and ClC-2, but not MLC1, have been found to colocalize in myelinated fiber tracts ^8^.

To gain insight into the biological processes affected by the GlialCAM cytoplasmic domain truncation, we conducted DEG analysis and found unique transcriptional changes. In GlialCAM ΔCT mice, the expression of genes associated with the neuroinflammation, including complement factor C4b and the serine peptidase inhibitor Serpina3n, were upregulated in both cortical and cerebellar cells. Serpina3n and C4b are markers of reactive astrocytes and oligodendrocytes ^30,31^. In the cerebellum, Serpina3n upregulation was restricted to Hepacam-enriched cell types, namely astrocytes and oligodendrocytes. In contrast, within the cortex, Serpina3n transcription was upregulated not only in astrocytes and oligodendrocytes, but also in some neuronal cells, suggesting that GlialCAM can differentially regulate inflammatory signaling in a region-specific manner. Serpina3n is involved in proteolytic pathways that regulate inflammation, immune responses, and cellular invasion during tissue remodeling ^32^. Inflammatory roles vary depending on its cellular origin. Astrocyte-derived expressed Serpina3n is associated with promoting neuroinflammation ^33^, while neuron-derived expressed Serpina3n has been shown to suppress neuroinflammatory responses ^34^. Consistent with this, in the GlialCAM ΔCT mice, Iba-1^+^ cells were increased only in the cerebellum, where Serpina3n upregulation is restricted to astrocytes and oligodendrocytes. In the cortex, however, where Serpina3n upregulation occurs in glial and neuronal cells, Iba-1^+^ cell numbers remained unchanged. These findings suggest that GlialCAM truncation drives inflammatory responses predominantly in the cerebellum, likely through astrocyte-derived Serpina3n. In GlialCAM ΔCT mice, Serpina3n upregulation was observed alongside elevated C4b transcription in both oligodendrocytes and astrocytes in the cortex and cerebellum. However, In the cerebellum of GlialCAM ΔCT mice, C4b upregulation was specifically restricted to oligodendrocytes. Several neurodegenerative and neuroinflammatory conditions, such as Alzheimer’s disease ^31^ and epileptic seizures ^34^, correlate with upregulation of C4b and Serpina3n in astrocyte and oligodendrocyte populations.

The ECM is highly dynamic, constantly undergoing remodeling to regulate various aspects of brain physiology. The ECM plays an essential role in modulating signaling pathways by influencing the distribution, activation, and presentation of ligands to their respective cellular receptors ^35^. Several genes associated with ECM organization, including Sparc, Fmod, Fbln5, and Adamts1, were upregulated in astrocytes and oligodendrocytes from the GlialCAM ΔCT cerebellum. Sparc, a Ca^2+^-dependent secreted glycoprotein, binds directly to collagen, modulating its assembly and crosslinking ^36^. Sparc also influences collagen deposition through the production and activation of Adamts1, a secreted metalloprotease ^37^. Conversely, Sparc overexpression has been linked to cell invasion through decreasing collagen IV levels and weakening the basement membrane ^38^. Its roles in promoting ECM disassembly and degrading ECM networks are regulated by Ca^2+^ ^39^. Some mutant mouse models of MLC display defects in intracellular Ca^2+^ homeostasis in response to osmotic stress ^40^. Furthermore, in the brain, chondroitin sulphate proteoglycans (CSPGs), which are known substrates for Adamts1, are upregulated in astrocytes in response to transforming growth factor β1 (TGFβ1) and epidermal growth factor (EGF) ^41^. While the protease activities of Adamts1 are essential for removing excess CSPGs to support optimal neurite outgrowth, excess Adamts1-mediated CSPG degradation may potentially promote ECM breakdown ^42^.

Fibromodulin (Fmod), a member of the leucine-rich proteoglycan family, was specifically upregulated in GlialCAM ΔCT cerebellar astrocytes. As a secreted protein, Fmod is expressed in different connective tissues, where it binds to collagen I and II, inhibiting fibril growth and resulting in thinner collagen fibrils ^43–45^. Additionally, Fmod has been identified as an activator of the complement system. Studies demonstrate that in human serum, Fmod interacts with C1q via its N-terminal tail to activate C1, subsequently triggering complement activation and C4b deposition ^46^. Fibromodulin has also been detected in the astrocyte secretome ^47^. However, its exact biological roles in astrocytes remain unknown. Furthermore, Fmod has been identified as a gene target regulated by FBLN5 in fibulin-5 overexpressing 3T3 fibroblasts, leading to reduced NFkB activity and elevated fibroblast apoptosis ^48^.

Furthermore, for the first time, our study demonstrates that selective truncation of the GlialCAM cytoplasmic tail induces specific deficits in coordination, hindlimb strength, and memory. In GlialCAM ΔCT mice, reduced hindlimb strength was evident from detailed gait analysis. Spatial and temporal parameters on the CatWalk system showed increased paw intensity values, with forelimbs exhibiting higher mean and peak intensities at maximal contact. This likely reflects a compensatory shift in body weight distribution to the forelimbs, minimizing load on the weaker hindlimbs. In addition, truncation of the GlialCAM cytosolic domain disrupted interlimb coordination ^49^. Interestingly, GlialCAM ΔCT mice displayed a slight but measurable increase in both Mlc1 transcript and protein levels. Therefore, the phenotypes observed in our truncation model are not attributable to MLC1 deficiency but likely reflect a loss of GlialCAM-specific functions, further distinguishing this model from MLC1 null phenotypes.

To our knowledge, this is the first proteomic study that systematically identifies interactions binding specifically to the cytoplasmic tail of GlialCAM in the cerebellum. These interactions appear to mediate intracellular signaling events, providing mechanistic insights into how GlialCAM extends its function beyond intercellular adhesion. Indeed, our work highlights the critical role for the GlialCAM cytoplasmic tail as a central signaling hub linking membrane adhesion complexes with key intracellular pathways, including MAPK signaling, cytoskeletal regulators and several intracellular effectors. Through these interactions, we propose that the cytosolic tail coordinates cellular processes essential for axon-glia interactions that govern maintaining white matter structural hemostasis. Indeed, recent studies have implicated dysfunction of the GlialCAM cytoplasmic tail in the pathogenesis of multiple sclerosis, a disorder characterized by neuroinflammation and white matter degeneration in the CNS ^50^. Specifically, immune responses against a highly conserved peptide sequence within the Epstein-Barr virus nuclear antigen 1 cross-react with the cytoplasmic tail of GlialCAM. It is interesting to speculate that defective signaling via the cytoplasmic tail promotes demyelination in multiple sclerosis patients similar to what we observe in GlialCAM ΔCT mutant mice.

## METHODS

### Animal model and genotyping

All experiments were performed in compliance with the Institutional Animal Care and Use Committee (IACUC) and The University of Texas MD Anderson Cancer Center Subcommittee on Animal Studies, both of which are AAALAC-accredited institutions. C-terminally truncated GlialCAM mice were generated using Cre/loxP technology. The C-terminal tail of GlialCAM is encoded by exons 5-7. To generate a C-terminal deletion variant of GlialCAM, a premature stop codon was inserted at the end of exon 4. To minimize potential dominant-negative effects, a cDNA mini-gene containing the coding sequences of exons 5-7, along with a polyA tail, was inserted upstream of exon 5. This mini-gene is flanked by loxP sites, allowing Cre-mediated deletion. Once deleted, transcription proceeds to the premature stop codon in exon 4, resulting in a C-terminally truncated GlialCAM variant (ConKO/GCAMΔCT). Nestin-Cre mice (stock 003771) and Rosa26-loxSTOPlox-YFP mice (stock 006148) were purchased from Jackson Laboratories.

### Cell culture and immunoblotting

Astrocytes were cultured as described previously ^51^. Briefly, cerebral cortices were dissected from neonatal brains (P1-P5) followed by removing the meninges. The cortical tissue was finely minced and collected by centrifuging at 1000 rpm in low-glucose Dulbecco’s Modified Eagle’s Medium (DMEM) supplemented with L-glutamine and sodium pyruvate media (10-014-CV, Corning). The pellet was resuspended in DMEM containing 150 U/ml collagenase (Type I, LS004196, Worthington Biochemical) and 40 µg/ml DNase I (D7291, Sigma-Aldrich) and digested for 30 minutes at 37°C. The resulting cell suspension was filtered through a 70-um nylon cell stainer into astrocytic growth medium (low-glucose DMEM with 10% bovine calf serum [BCS; SH30072.03, Hyclone] complemented with 1 U/ml penicillin-streptomycin [MT30004CI, Corning]). Finally, the cells were seeded into a T-75 tissue culture flask precoated with laminin. Astrocytes isolated from N-Cre^-^:GCAM^ConKI/+^ and N-Cre^-^:GCAM^ConKI/ConKI^ P2 pups were dissociated, and the cultured cells were infected with either Adeno (control, UIOWA, Ad5CMVempty) or Adeno-Cre viruses (UIOWA, Ad5CMVCre) in serum free media. 48 hours after transfection cells were collected in DNA lysis buffer for genomic PCR analysis. Mice brains were dissected and homogenized in lysis buffer (RIPA buffer 89901, Thermo Fisher Scientific) containing proteinase and phosphatase inhibitors (A32955, A32957, Thermo Fisher Scientific). Protein extracts were separated on 7% or 10% SDS-PAGE gels. Primary antibodies and dilutions used for immunoblotting included the following: Rabbit anti-Hepacam (Abcam, ab300571; 1:1000); Rabbit anti-CSK (Cell Signaling, 4980; 1:1000); Rabbit anti-PDK1 (Cell Signaling, 3062; 1:1000); Rabbit anti-MMP14 (Abcam, ab51074; 1:1000), Rabbit anti-SOX2 (Cell Signaling, 2748; 1:1000); Mouse anti-GAPDH (60004, Protein technology; 1:50,000). horseradish peroxidase–conjugated secondary antibody (goat anti-rabbit, Bio-Rad, 1705046; 1:5000) or (goat anti-mouse, Bio-Rad, 1705047; 1:5000) was used as a secondary antibody. The signal was detected with an enhanced chemiluminescence reagent (PerkinElmer, NEL 104001EA).

### Cell surface biotinylation

Mouse astrocytes were kept on ice and washed three times with ice-cold PBS before proceeding to treating them with biotin. Cells were then incubated with 10 mM Sulfo-NHS-Biotin (Thermo Scientific) in PBS+Ca²⁺ on ice for 30 minutes, followed by three washes with PBS with Ca²⁺ containing 100 mM glycine. Next, the cells were resuspended in 250 μl of lysis buffer (composed of 1% Triton X-100, 25 mM HEPES, 150 mM NaCl, 10 mM MgCl₂, 1 mM EDTA, a protease inhibitor mixture from Roche Diagnostics, and 2% glycerol) and lysed by gentle rotation on a spinning wheel at 4 °C for 30 minutes. The soluble fraction was incubated with 50 μl of streptavidin agarose beads (Thermo Scientific) for 1 hour at 4 °C. Following this, the beads were washed three times with lysis buffer and resuspended in 35 μl of Laemmli buffer. Biotinylated proteins were subsequently detected by Western blot, as described earlier.

### immunohistochemistry and immunofluorescence

Adult mice brains were harvested and fixed overnight in cold 4% paraformaldehyde (PFA) in PBS. Fixed brains were washed three times in PBS and embedded in paraffin. Formalin-fixed, paraffin-embedded brain tissues were deparaffinized at 65C and rehydrated in decreasing concentrations of alcohol. Heat-induced antigen retrieval was performed in the presence of proteinase K (20mg/ml) in TE buffer (10mM Tris, 1mM EDTA, pH=9) for 8 minutes. Sections were permeabilized with 0.2% Triton X-100 in PBS and blocked in 10% normal horse serum for 1 h at room temperature. Tissue sections were incubated overnight at 4C with the primary antibody rabbit anti-MBP (Abcam, ab40390, 1:1000) or chicken anti-Neurofilament (Neuromics, CH22104, 1:5000) diluted in 1% BSA. The following day, sections were incubated with a biotinylated secondary antibody (horse ant-rabbit, BA-110, 1:400) in blocking buffer for 1 h at room temperature. After washing with PBS, the sections were incubated with ABC reagents (PK-4000, Vector Laboratories) for 45 min at room temperature. Signal visualization was achieved using ImmPACT DAB substrate (SK-4105, Vector Laboratories) for 5 minutes, followed by multiple washes in double-distilled water. Counterstaining was performed with hematoxylin for 45 seconds, followed by rinsing in cold tap water. Sections were then dehydrated in increasing concentrations of alcohol and mounted with mounting media. Images were acquired using Olympus BX43 light microscope. Alternatively, fixed brains were embedded in 3% agarose and sectioned at 100 μm on a vibratome.

For preparing cryosections, fixed brains were incubated in 30% sucrose before being embedded in OCT. After heating sections at 37C for 10 minutes, they were blocked in 10% normal donkey serum with 1% BSA and 0.3% Triton X-100 for 1 hour at room temperature. Tissues were then incubated with the primary antibodies diluted in 1% normal donkey serum, 0.1% BSA, and 0.3 Triton x-100. Primary antibodies were as follow: goat-anti CD31 (R&D Systems, AF3628, 1:200); rat anti-CD31 (BD Pharmingen, 553370, 1:100); chicken anti-GFAP (Novus, 05198, 1:1000,); rabbit anti-Iba-1 (FUJIFILM, 1919741, 1:200), goat anti-Iba-1 (Abcam, ab5076, 1:500); rabbit anti-Hepacam (599999; 1:200); rat anti-MBP (Abcam, ab7349, 1:1000); goat anti-Serpina3n (R&D Systems, AF4709; 1:100); goat anti-Olig2 (R&D Systems, AF2418; 1:200); chicken anti-GFP (Abcam, ab13970, 1:500); rabbit anti-GFP (Novus Biological, NB600308, 1:500); rabbit ant-PDGFRα (Abcam, ab203491, 1:200); and rabbit anti-MLC1 (Invitrogen, PA5-41042, 1:100). After washing with PBS, sections were incubated for 45 minutes at room temperature with fluorochrome-conjugated secondary antibodies (Jackson ImmunoResearch, 1:500),.

### BBB permeability analysis

Transient focal ischemia was induced for 45 minutes by suture occlusion of the middle cerebral artery (MCAO) in male mice anaesthetized using 1.5% isoflurane, 70% N2O and 30% O2. Ischemia was induced by introducing a coated filament (702223; Doccol) from the external carotid artery into the internal carotid artery and advancing it into the circle of Willis to the branching point of the left MCA, thereby occluding the middle cerebral artery. Achievement of ischemia was confirmed by monitoring regional cerebral blood flow (CBF) in the left middle cerebral artery. CBF was monitored through a disposable microtip fiber optic probe (diameter 0.5 mm) connected through a laser doppler monitor (moorVMS-LDF). The microtip was attached to the skull of the mouse through glue. Animals that did not show a CBF reduction of at least 80% were excluded from the experimental group. Rectal and temporalis muscle temperature was maintained at 37°C with a thermostatically controlled heating pad. All surgical procedures were performed under an operating stereomicroscope. After 4 hours, mice were injected retro-orbitally with Alexa Fluor 488-cadaverine.BBB integrity was assessed through retro-orbital delivery of Alexa Fluor 488-cadaverine (640 Da; Invitrogen, A30676) at a dose of 6 µg per gram of body weight. Following a 2-hour circulation period, brains were collected after cardiac perfusion with 4% paraformaldehyde PFA. Brain sections were processed as described earlier.

### Motor coordination and cognitive assays

Beam Walk: The beam walk task evaluates motor coordination, balance, and sensory and proprioceptive sensitivity in mice. Mice were trained on three consecutive days to cross a 12 mm-wide rectangular beam (‘wide flat’), followed by a 6 mm-wide rectangular beam (‘narrow flat’) and a 6 mm-diameter cylindrical beam (‘round’). All three beams are 85 cm in length and placed at an approximate 15-degree uphill incline. The time taken to cross, and the number of errors (paw slips, inversions) were determined by investigators blinded to treatment. Mice that fell from the beam or failed to complete the task within 30 seconds (wide flat), 45 seconds (narrow flat) or 60 seconds (round) were assigned the cutoff time of 30, 45 or 60 seconds respectively.

CatWalk: Analysis of gait patterns was performed using the Catwalk® XT 10.6 system (Noldus Information Technology, Leesburg VA) as previously described ^52^. All habituation and testing were performed in the light cycle between 12:00 and 16:00 h. Four compliant runs were captured per mouse (run duration between 0.5 and 10 s, maximum variation in run speed ≤60%).

Grip Strength: The grip strength test was performed as previously described ^53^. Hindlimb grip strength was measured using a Bioseb grip strength meter. All habituation and testing were performed in the light cycle between 12:00 and 16:00 h. Mice were scruffed, and once both hind paws gripped the wire mesh, the mouse was slowly pulled away from the grip strength meter. The max pull force in grams was recorded.

Novel object place recognition test (NOPRT): The NOPRT for short-term memory and place recognition was performed as previously described ^54^ with some modification. During training, mice were introduced to two identical objects placed on one side of a rectangular arena for 10min. After 24 h, mice were returned to the arena containing one familiar object at the same location as in the training session, and one novel object (Rubik’s cube) placed in a novel location. Interaction times with each of the objects were recorded for 10 min and analyzed with EthoVision XT 10.1 video tracking software (Noldus Information Technology Inc., Leesburg, VA). The discrimination index was calculated as (TNovel − TFamiliar)/ (TNovel + TFamiliar). Finally, to assess hindlimb clasping, each mouse was suspended by the base of its tail for 30 seconds above a platform free of surrounding objects.

### Single-cell RNA-seq sample preparation

Single-cell RNA sequencing was performed on fixed cortical and cerebellar cells isolated from two 1-year old WT and GCAM ΔCT mice using the Chromium Fixed RNA Profiling workflow from 10x Genomics. Following tissue dissociation and fixation, single-cell suspensions were prepared in accordance with the manufacturer’s guidelines. To enable multiplexed analysis, samples were pooled and uniquely barcoded. Whole-transcriptome probe pairs were hybridized to the fixed cells overnight to specifically capture target RNAs. Barcoded Gel Beads-in-Emulsion (GEMs) were then generated using the Chromium Next GEM platform to achieve cell- and sample-specific indexing. cDNA libraries were subsequently constructed and sequenced on an Illumina platform. The resulting data were demultiplexed and processed to generate single-cell gene expression profiles.

### Xenium sample preparation

For spatial profiling using Xenium, 5-μm-thick formalin-fixed paraffin-embedded (FFPE) mouse brain tissues were mounted onto a Xenium slide (10x Genomics). The sample preparation followed the procedures detailed in the 10x Genomics User Guides (CG000578, CG000580, CG000749, and CG000584). Briefly, tissue sections underwent deparaffinize, rehydration through an ethanol gradient, and de-crosslinking. Next, overnight hybridization was performed using both the pre-designed Mouse Brain Panel (10x Genomics) and the add-on custom panel. Subsequent steps included probe ligation and enzymatic amplification to generate rolling circle amplification (RCA) products. Prior to imaging, cell boundaries were segmented, autofluorescence was reduced, and nuclei were stained. Finally, the samples were loaded onto the Xenium Analyzer instrument, where regions of interest were selected based on DAPI-stained images captured by the instrument.

### Data processing and Quality Control

ScRNA-seq data were processed using Cell Ranger (v5.0) from 10x Genomics for demultiplexing, alignment, and gene expression quantification. To further remove background counts from ambient RNA and barcode swapping, CellBender (v0.3.0) was applied to the gene-barcode matrices. Although whole brain samples were used in our Xenium data, only cortex and cerebellum cell IDs were extracted from samples using the Explorer software (10X Genomics) and used for downstream analysis.

Downstream analysis of scRNA-seq and Xenium data was performed using the Seurat R package (v4.2.1) on RStudio 4.4.1. Cells were filtered based on quality control metrics, including the number of detected genes, total RNA counts, and mitochondrial gene content. After normalization and scaling, Seurat object for each sample were merged and dimensionality reduction was performed using principal component analysis (PCA), followed by uniform manifold approximation and projection (UMAP) for visualization. Clusters were identified with graph-based clustering, and using the FindConservedMarkers function, conserved marker genes were identified across conditions. Canonical marker genes were used to assign cell types. Bar plots were used to visualize percentage of cell types in each sample. Dot plots were used to visualize transcript percent expression and average expression of gene markers per cell type. Annotated cell IDs were exported from R and imported into the 10x genomics Xenium software to visualize cell clusters.

Differential gene expression analysis of scRNA-seq data between GlialCAM ΔCT and N-Cre samples was performed using Seurat’s FindMarkers function (v4.2.1) with the Wilcoxon rank-sum test. Genes were required to be expressed in at least 10% of cells in either group (min.pct = 0.1). Heatmaps of differentially expressed genes were generated with heatmap, displaying scaled expression values.

For the xenium data, heatmap visualizations were generated using the DoHeatmap function from the **Seurat** R package to display the expression patterns of selected genes across defined cell groupings. Cells were grouped according to their condition (i.e., GlialCAM ΔCT, N-Cre). Expression values were scaled by default within Seurat to highlight relative differences across groups.

To assess how annotated clusters map back to tissue architecture, spatial images were created using the ImageDimPlot function from the Seurat package which plots spatial distribution of cells based on their tissue coordinates (extracted from Xenium object metadata). Cells are pseudocolored according to their cluster annotation.

To generate a spatial density map, the 2D spatial coordinates (X, Y) of each cell (extracted from Xenium object metadata) were binned (bin size = 100) into discrete grid regions (x_bin, y_bin), and the number of cells per bin was counted to compute a **localized cell density**. The resulting binned count data was visualized using the plot_ly function from the **Plotly package** in R (version 4.11.0), with each bin represented as a square marker. This visualization allows for intuitive interpretation of spatial enrichment patterns, revealing localized cell distributions differences between GCAM ΔCT and N-Cre cortex.

### Revers-Phase Protein Array sample preparation

Astrocytes isolated from P1 pups were seeded on a laminin coated dish and incubated till reaching confluent. Adherent astrocytes were washed with ice-cold PBS and lysed in RPPA lysis buffer (1% Triton X-100, 50 mM Hepes (pH 7.4), 150 mM NaCl, 1.5 mM MgCl₂, 1 mM EGTA, 100 mM NaF, 10 mM sodium pyruvate, 1 mM Na₃VO₄, 10% glycerol, and a cocktail of protease and phosphatase inhibitors (Roche Diagnostics)). Lysates were denatured by heating at 95°C for 5 minutes in 4× SDS sample buffer without β-mercaptoethanol. Samples were processed and probed with 499 antibodies at the functional proteomics RPPA core facility at MD Anderson. The RPPA data were normalized for protein loading.

### Expression and purification of GST-GlialCAM C-terminal tail

*E. coli* BL21 cells were transformed with the pGEX-6P-1 vector harboring the entire C-terminal tail of *Hepacam* (474bp) fused to a GST tag. An empty vector (EV) served as the control. These constructs were purchased from GenScript. Recombinant protein expression in *E. coli* BL21 cells was induced with 0.5 mM Isopropyl β-D-1-thiogalactopyranoside (IPTG, Sigma #16758) for 3.5 hours. Cells were lysed in Lysis buffer (50mM Tris pH=7.4, 150mM NaCl, 5mM MgCl2, 1mM DTT, 1mM PMSF, 1%Triton x-100, protease inhibitor). The clarified lysate was incubated with 1ml of pre-washed Glutathione Sepharose 4B beads (Cytiva #17075601) for one hour at 4 °C. Beads were collected by centrifugation at 1000rpm for 2minutes, washed with lysis buffer containing 0.5% Triton x-100, and resuspended in GST freezing buffer (wash buffer supplemented with 10% autoclaved glycerol). The bead-protein complex was stored at −80 °C until use in immunoprecipitation (IP) experiment.

### GST-GlialCAM C-terminal tail Immunoprecipitation

The whole cerebellum from a 7-month-olde mouse was homogenized in the homogenizing buffer (50mM Tris buffer pH 8.0, 2 mM MgCl2, 1mM EDTA, and a Complete protease and phosphatase inhibitor tablets). Crude membranes were collected by centrifugation and solubilized in IP buffer (50 mM Tris-Cl pH 7.4, 150 mM NaCl, 1%TritonX-100, and a Complete protease and phosphatase inhibitor tablets) for 2h at 4°C. The supernatant was incubated with 60ug of the glutathione bead-GST-GC-Cyto/EV complex for 3h at 4°C. in parallel, an equivalent amount of beads were incubated only with IP buffer served as an additional control. Beads were washed 5 times with IP buffer and eluted with 50ul 2x SDS sample buffer, followed by heating at 37°C for 10min. The IP samples were resolved by SDS-PAGE and visualized by Coomassie staining.

### Mass Spectrometry sample preparation and analysis

Excised gel bands were thoroughly de-stained and dehydrated using acetonitrile, followed by trypsin (V5113, Promega Corporation, Madison, WI) digestion in 50 mM NH4HCO3 at 37 °C overnight. The resulting peptides were extracted with acetonitrile and dried under vacuum. The vacuum-dried tryptic peptides were dissolved in solvent A (0.1% formic acid in water) and directly loaded onto a nano reverse-phase C18 column (75 μm ID × 25 cm L, 1.7 μm particle size). Peptides were separated using a linear gradient of solvent A (0.1% formic acid in water) and solvent B (0.1% formic acid in 80% acetonitrile). The gradient was as follows: 2% to 8% solvent B over 2 minutes, 8% to 38% solvent B over 45 minutes, 38% to 100% solvent B over 1 minutes, and held at 100% solvent B for 11 minutes. The eluate was electrosprayed into an Orbitrap Astral Mass Spectrometer (Thermo Fisher Scientific, Waltham, MA) operating in positive ion mode, with an electrospray voltage of 2 kV and an ion transfer tube temperature of 280°C. Data were acquired in DDA mode with a fixed cycle time of 3 seconds. Full MS scans were performed in the range of 350 to 1350 m/z at a resolution of 240,000, with an AGC target of 300% and a maximum injection time of 100 ms. Precursor ion selection width was set to 2 Th. Peptide fragmentation was carried out using HCD with a normalized collision energy of 25%. Fragment ion scans were recorded by Orbitrap with a resolution of 60,000, AGC target of 300%, and a maximum injection time of 100 ms.

For data analysis, the MS/MS spectra were processed using Proteome Discoverer 2.4 (Thermo Fisher Scientific) with a mouse protein sequence database from UniProt (17184 entries, downloaded November 2023). Peptide searching was performed with Sequest HT using the following parameters: allowing up to two missed cleavages, a precursor mass tolerance of 10 ppm, and a product ion tolerance of 0.6 Da. Oxidation (M) and Acetyl (N-Terminus) were set as variable modifications.

## Supplemental Figure Legends

**Supplemental Figure 1.**
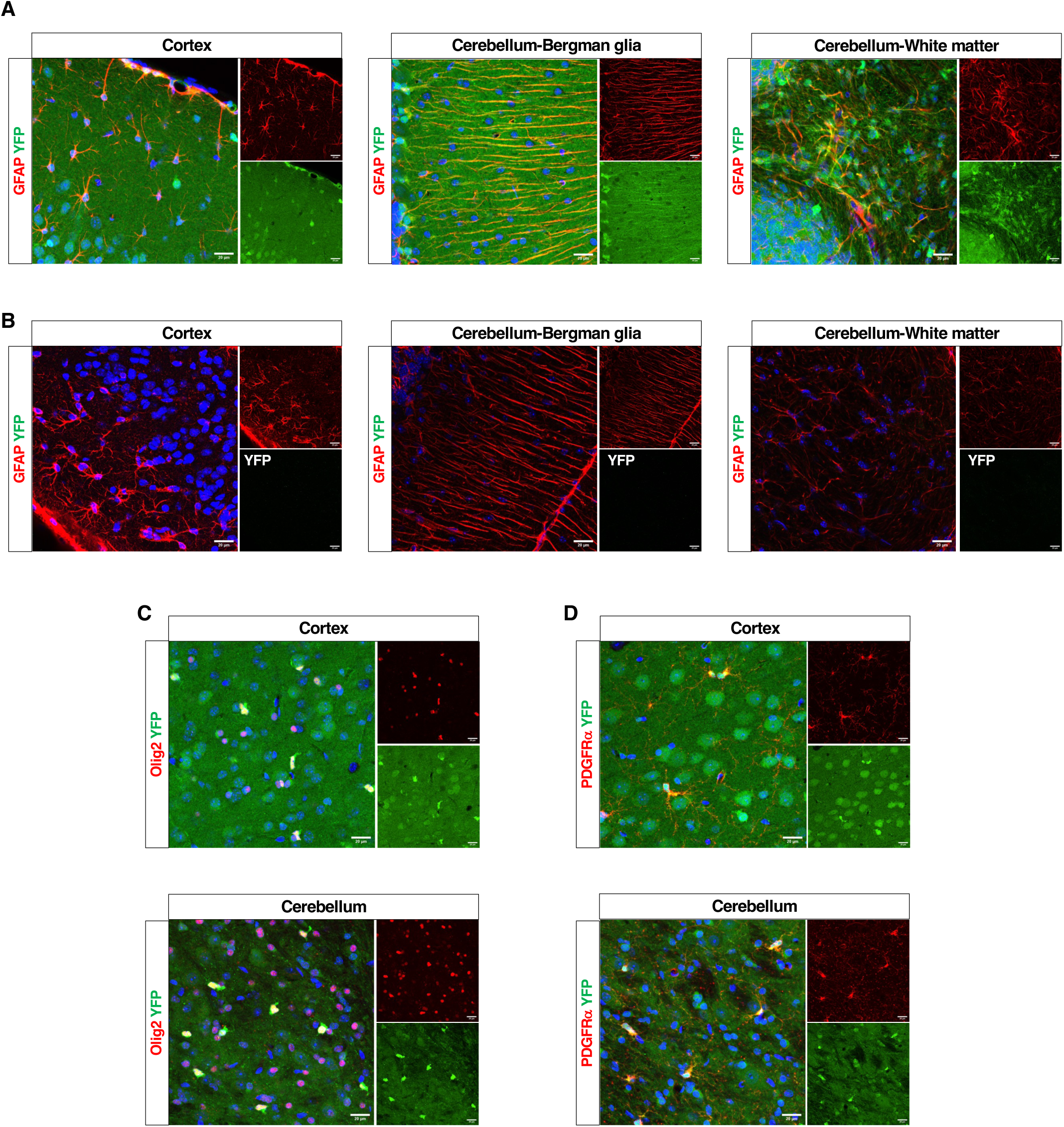
Analysis of Nestin-Cre mediated recombination in adult mice glial cells. **(A-D);** Immunofluorescence staining of cortex and cerebellum sections from adult N- Cre/+:Rosa26-loxSTOPlox-YFP/+ (A, C, and D), and Rosa26-loxSTOPlox-YFP/+ mice. Sections were stained with anti-GFP (green) together with anti-GFAP (red) (A, B), anti-Olig2 (red) (B), or anti PDGFRα (red) (C) antibodies. Nuclei are counterstained with DAPI. Scale bars, 20 μm.

**Supplemental Figure 2.**
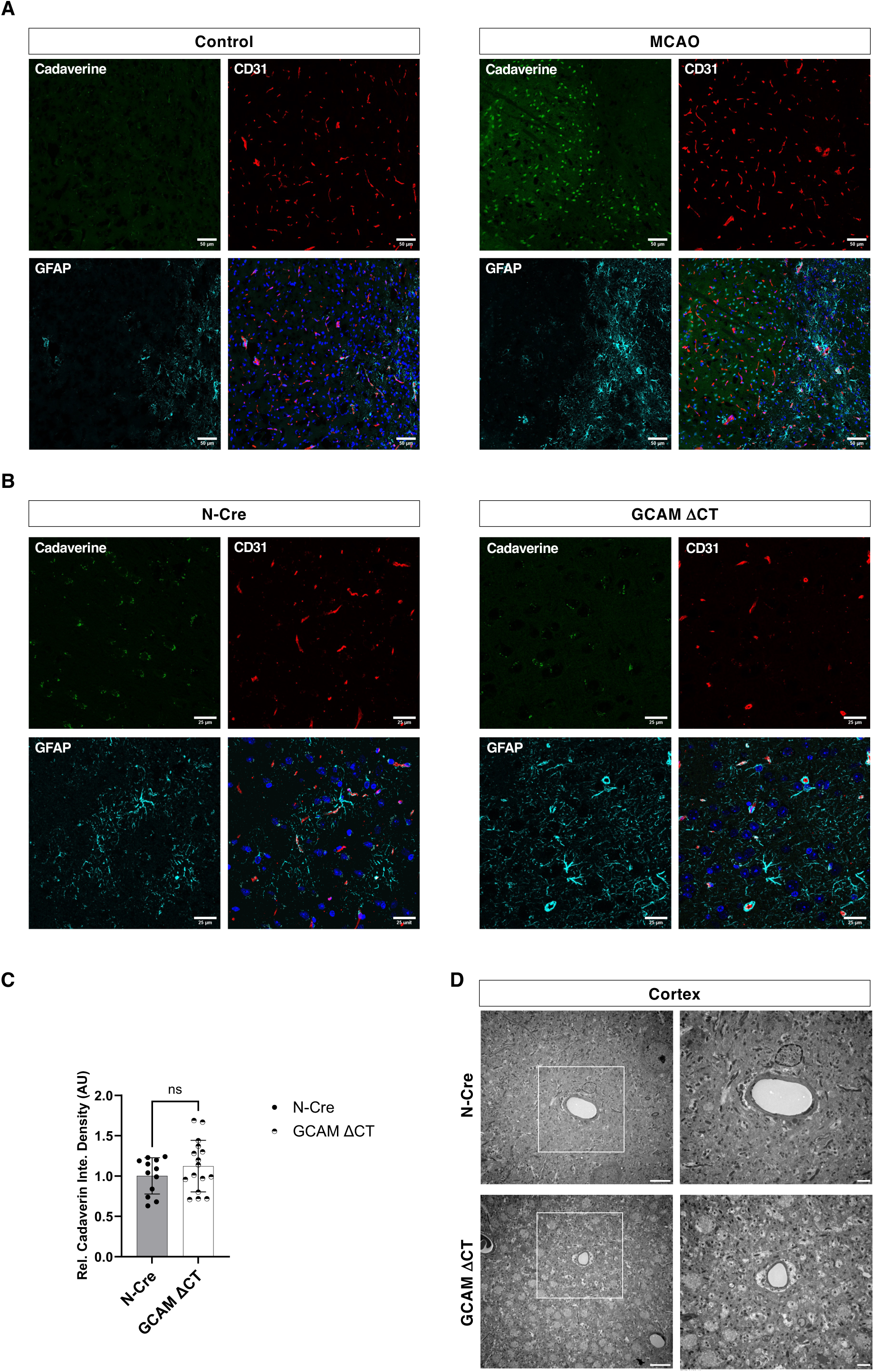
The blood brain barrier remains intact in GlialCAM C-terminally truncated mice. **(A);** Immunofluorescence staining of brain sections from an adult wild type mouse subjected to 45 minutes of MCAO. Mice received intra-orbital injection of 488-cadaverine 4 hours later and were perfused within 2 hours. Sections were stained with anti-CD31 (red), and anti-GFAP (cyan), with nuclei counterstained using DAPI. Scale bars, 50 μM. **(B);** Immunofluorescence staining with anti-CD31 (red), and anti-GFAP (cyan) was performed on brain sections from 12-month-old control (N-Cre) and GlialCAM (GCAM) ΔCT mice, collected 2 hours after intra-orbital injection of 488-cadaverine. Nuclei are counterstained with DAPI. Scale bars, 25 μM. **(C);** Quantification of extravasated cadaverine across 3 biological replicates of mice represented in (B). **(D);** Transmission electron microscopy analysis of brain sections from the cortex of 12-month-old Control (N-Cre) and GlialCAM (GCAM) ΔCT mice. Boxed areas in the left panels are shown at higher magnification in the right panels. Scale bars, 6 and 2 μM. All data are shown as mean ± SD. Statistical significance was determined by two-way ANOVA.

**Supplemental Figure 3.**
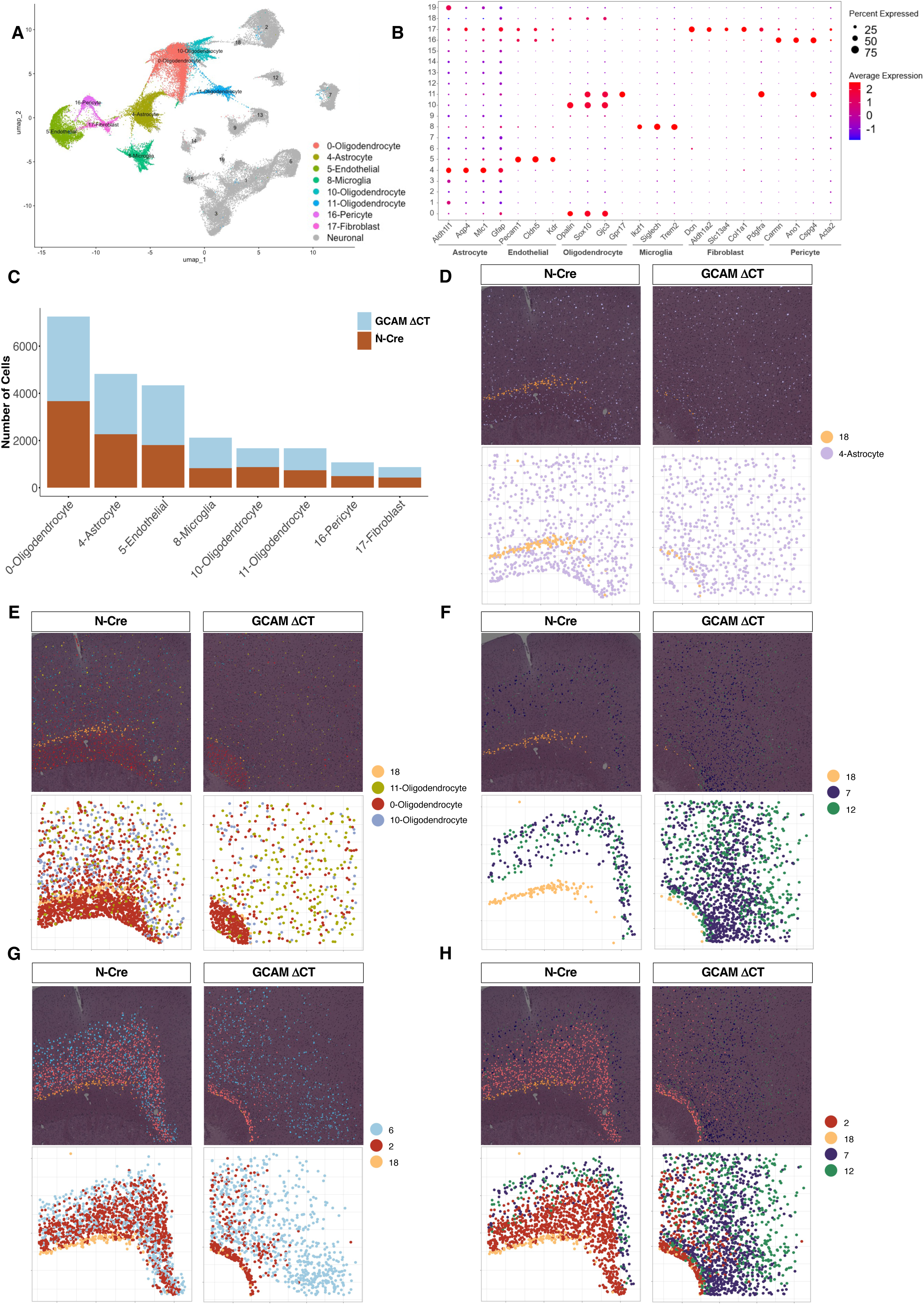
In situ spatial transcriptomics analysis of the cortex and corpus callosum regions in GlialCAM ΔCT and control mouse brains. **(A);** UMAP visualization of 20 clusters identified in the cortex and corpus callosum regions of control (N-Cre^+^) and GCAM ΔCT mice. **(B);** Dot plot depicting the expression levels of representative cell-type-enriched marker genes across all clusters. **(C);** Bar graph illustrating the percentage of non-neuronal cell types across all cells identified within the cortex and corpus callosum of control (N-Cre^+^) and GCAM ΔCT mice. **(D-G);** Special transcriptomic data from the frontal cortex of 12-month-old control (N- Cre^+^) and GCAM ΔCT mouse brains illustrating the spatial distribution of cells within the astrocyte cluster (D), oligodendrocyte clusters (E), neuronal layers V (clusters 7 and 12) (F), VI (cluster 2) (G), and V/VI (cluster 6) (G). **(H);** Highlights the boundary between layer V (cluster 7 and 12) and VI (cluster 2). In all panels, cluster 18 (layer 6b) is shown to distinguish cortex from the hippocampus region.

**Supplemental Figure 4.**
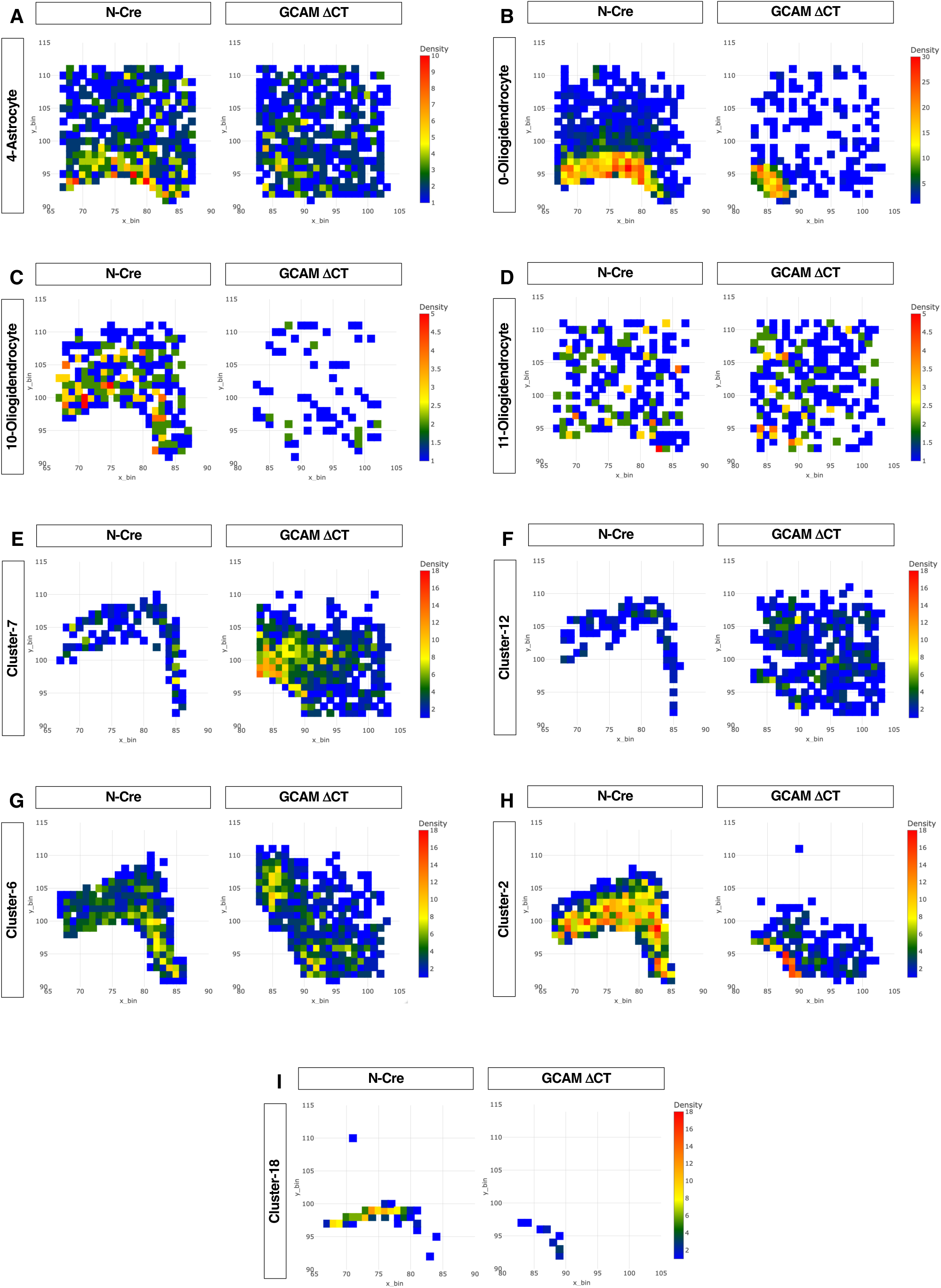
Spatial cell density maps of the frontal cortex in GlialCAM ΔCT and control mice. **(A-I);** Special transcriptomic data from the frontal cortex of 12-month-old control (N-Cre^+^) and GCAM ΔCT mouse brains illustrating the cell density distribution of cells within the 4-Astrocyte (A), 0-Oligodendrocyte (B), 10-Oligodendrocyte (C), 11-Oligodendrocyte (D), neuronal layers V (clusters 7 (E) and 12 (F)), V/VI (cluster 6) (G), VI (cluster 2) (H), and layer 6b (cluster-18) (I).

**Supplemental Figure 5.**
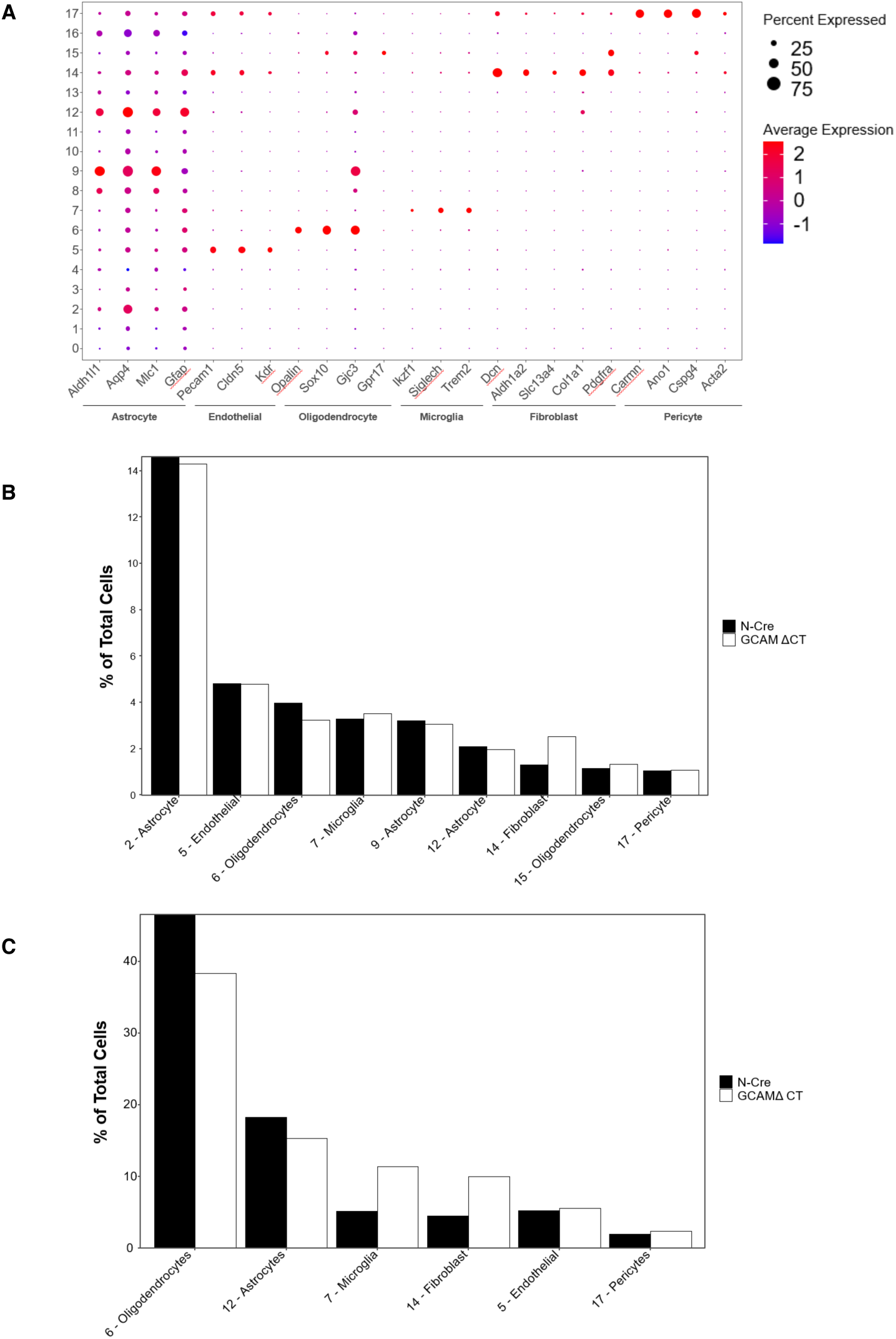
Cell type identification and composition reveled by spatial transcriptomic data of the cerebellum region of GlialCAM ΔCT and control mice. **(A)**; Dot plot depicting the expression levels of representative cell-type-enriched marker genes across all clusters. **(B)**; Bar graph illustrating the percentage of non-neuronal cell types across all cells identified within the whole cerebellum and white matter **(C)** region of control and GlialCAM (GCAM) ΔCT mice.

**Supplemental Figure 6.**
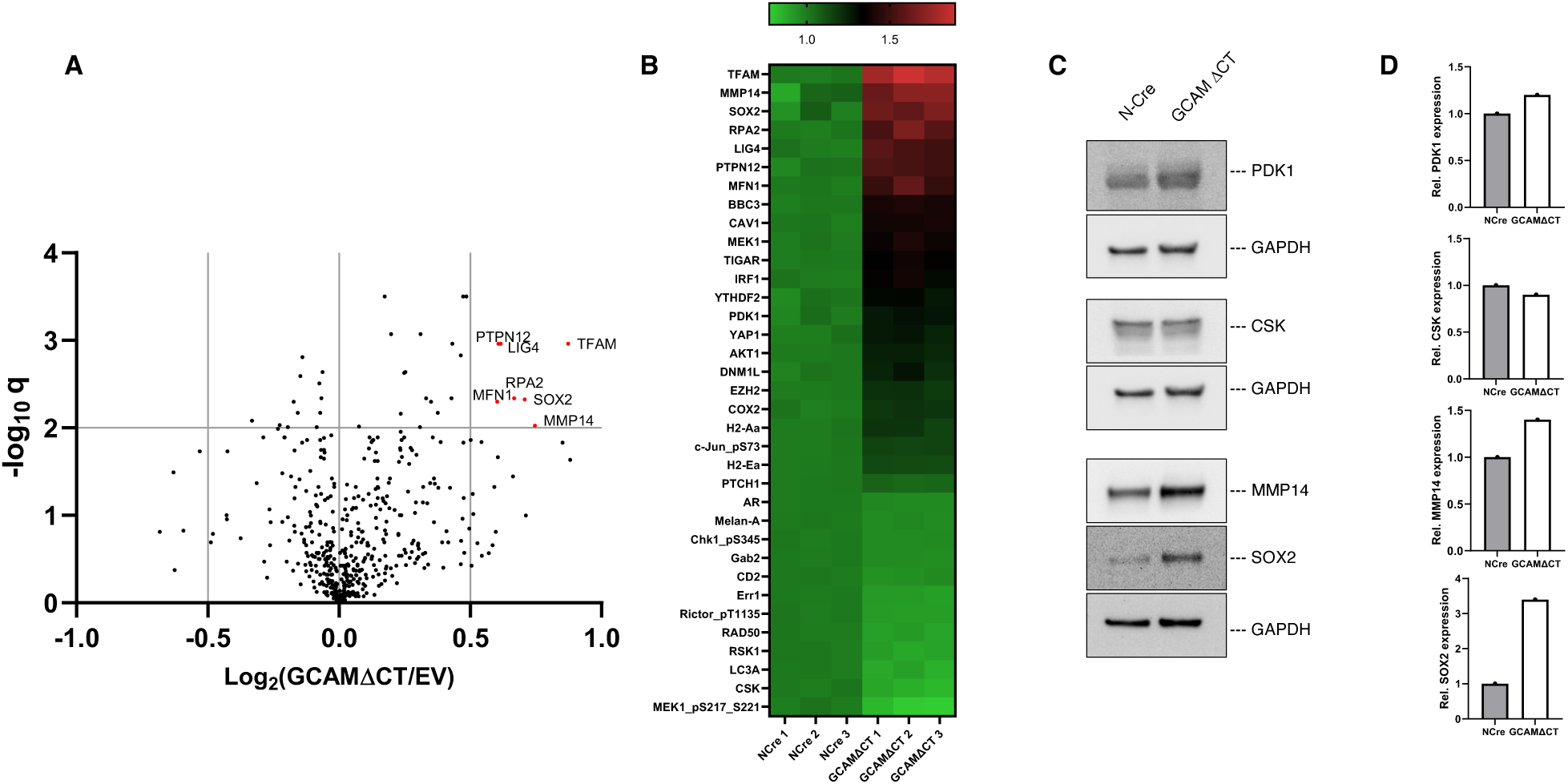
GlialCAM ΔCT mice displayed pronounced astrogliosis with upregulated MLC1 expression in astrocytes and increased Iba-1 immunoreactivity in the cortex and cerebellar white matter. **(A-C);** Immunofluorescence staining was performed on cerebellum (A, C) and cortex (B) sections from 12-month-old control and GlialCAM (GCAM) ΔCT mice. Sections were stained with anti-CD31 (green), anti-Iba-1 (red), and anti-GFAP (cyan) (A and B). Additional staining was performed with anti-CD31 (green), anti-MLC1 (red), and anti-GFAP (cyan) on cerebellum sections (C). Nuclei are counterstained with DAPI. Scale bars, 50 and 25 μM.

